# The innate immune systems of malacostracan crustaceans exhibit both conserved and evolutionarily distinct components

**DOI:** 10.1101/091835

**Authors:** Alvina G. Lai, A. Aziz Aboobaker

## Abstract

Growing demands for aquatic sources of animal proteins have attracted significant investments in aquaculture research in recent years. The crustacean aquaculture industry has undergone substantial growth to accommodate a rising global demand, however such large-scale production is susceptible to pathogen-mediated destruction. It is clear that a thorough understanding of the crustacean innate immune system is imperative for future research into combating current and future pathogens of the main food crop species. Through a comparative genomics approach utilising extant data from 55 species, we describe the innate immune system of crustaceans from the Malacostraca class. We identify 7407 malacostracan genes from 39 gene families implicated in different aspects of host defence and demonstrate dynamic evolution of innate immunity components within this group. Malacostracans have achieved flexibility in recognising infectious agents through divergent evolution and expansion of pathogen recognition receptors genes. Antiviral RNAi, Toll and JAK-STAT signal transduction pathways have remained conserved within Malacostraca, although the Imd pathway appears to lack several key components. Immune effectors such as the antimicrobial peptides (AMPs) have unique evolutionary profiles, with many malacostracan AMPs not found in other arthropod groups. Lastly, we describe four putative novel immune gene families, characterised by distinct protein domains, potentially representing important evolutionary novelties of the malacostracan immune system.

## Introduction

The global human population is projected to escalate to 9.1 billion by 2050 (Fischer et al., 2009). With an increasing food consumption per capita and changing demands for animal proteins (Kearney, 2010), there is a dire need for sustainable sources to avoid further degradation of the environment. It has been suggested that much of this may come from invertebrate sources including insects (Van Huis et al., 2013), but clearly crustaceans currently represent a source of protein that is more culturally palatable in Europe and North America. Crustaceans already represent a significant portion of marine aquaculture produce, with the predicted annual production exceeding 10 million tonnes and sales of $40 billion (Stentiford et al., 2012) that will continue to increase. The expansion of farmed crustaceans is not without major issues. It is estimated that up to 40% ($3 billion) of just shrimp production alone can be lost each year due to disease outbreaks (Stentiford et al., 2012). Pathogens and diseases impacting crustaceans have been recently extensively reviewed (Stentiford, 2012a; Stentiford et al., 2012b; Bondad-Reantaso et al., 2012; Flegel, 2012; Lightner et al., 2012; Shields et al., 2012). Some of the most common diseases in decapod crustaceans are white spot disease caused by the white spot syndrome virus (WSSV) in penaeid shrimps (Lightner et al., 2012), yellow head disease caused by the Yellow head Virus (Flegel 1997; Prapavorarat et al. 2010; Junkunlo et al. 2012), Taura syndrome caused by the Taura Syndrome Virus (Lightner et al. 1995; Stentiford et al. 2009), fungal diseases in the Dungeness crab, ***Cancer magister*** (Unestam 1973; Fisher and Wickham 1977; Armstrong et al. 1976), infections by the parasitic dinoflagellate ***Hematodinium*** sp. in crabs (Stentiford et al., 2003), the ***Panulirus argus*** virus 1 (PaV1) infection in lobsters (Behringer et al., 2011) and bacterial diseases caused by ***Vibrio*** or ***Aeromonas*** (Wang 2011). There is broad agreement that without new interventions and better understanding of pathology and immune responses, current best practices for crustacean aquaculture cannot be improved. The use of antibiotics and chemical treatments for disease control in aquaculture is undesirable due to long-term economic and environmental ramifications (Capone et al., 1996; Collier et al., 1998; Grant et al., 1998). Therefore, approaches that harness and aid the crustacean innate defence mechanism should be exploited to limit and prevent diseases, and therefore crop loss. For example, assays for the measurement of innate immune activity could provide early warnings for the presence of potential pathogens within closed aquaculture systems.

Systematic and cross-species characterisation of the crustacean immune system has not been performed, despite it being essential for the field to progress (Hauton, 2012). Previous comparisons amongst sequenced arthropod genomes of insects, chelicerates, the myriapod ***Strigamia maritima***, the branchiopod ***Daphnia pulex*** and the amphipod ***Parhyale hawaiensis*** have recently revealed signatures of conservation and diversity in innate immunity components across arthropod phyla (McTaggart et al., 2009; Palmer et al., 2015; Kao et al., 2016). However, not much is known about the evolutionary events that define the immune system in malacostracans, or within the order Decapoda that includes crop species. The radiation of Pancrustacea (hexapods and crustaceans) has been estimated to be between ~540 to ~666 million years ago (mya) (Pisani et al. 2004; Rota-Stabelli et al., 2013) while the split of Branchiopoda from Malacostraca was estimated at 614 mya (Pisani et al. 2004). Others have made estimates of similar divergence times based on crustacean hemocyanins (Hagner-Holler et al., 2005). Given the large evolutionary time scales involved, many lineage specific changes in immune system components within the Malocostraca may have occurred and using only the branchiopod ***D. pulex*** and a single malacostracan ***P. hawaiensis*** to define immune regulation is unlikely to provide either a rich or accurate picture. Ultimately this will require both comparative and functional genomics approaches to effectively understand and exploit the immune system. Due to potential importance of crustacean food sources, such studies are of high impact and urgency. Currently, the lack of a comprehensive comparative genomics study of immunity with the Malacostraca means that a clear staging point for underpinning this work is lacking.

Here, we address this major deficit by performing an in depth comparative study amongst the broader Malacostraca, including extant data from the order Decapoda that includes all the major food crop species (fig. 1). A large number of relatively recent independent studies have started to generate publically deposited large transcriptomic data sets from food crop species and other related malacostracan species providing ample raw data for our study (Christie et al., 2014; Ghaffari et al., 2014; Sun et al., 2015; Li et al. 2015; Meyer et al. 2015; Stahl et al., 2015; Verbruggen et al. 2015; Truebano et al. 2016; Kim et al. 2016; Havird et al. 2016; Tan et al., 2016; and complete set of references in supplementary table 1).

**Figure 1.**
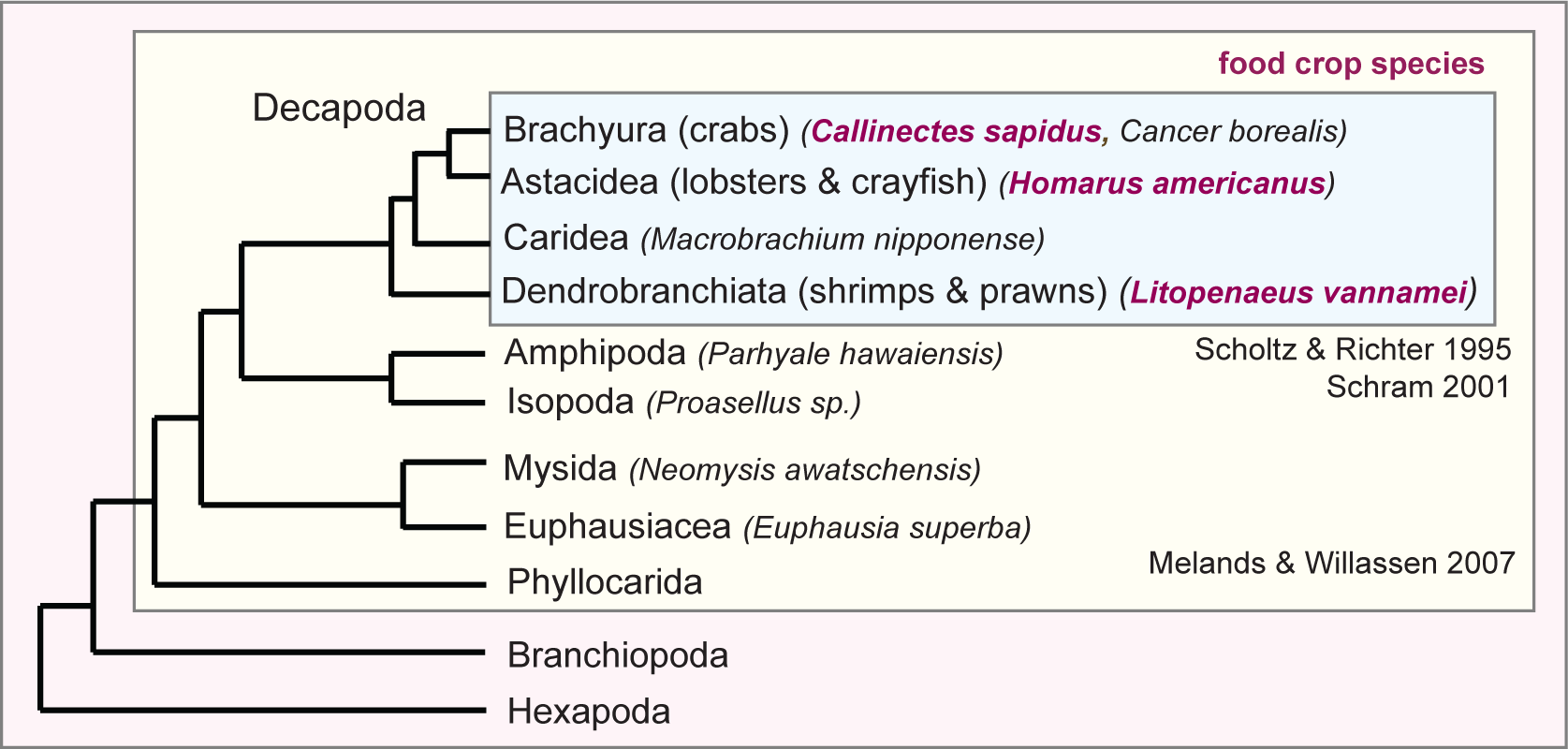
Phylogenetic relationship of Malacostraca. Malacostraca is shown within the Pancrustacea clade. Malacostraca tree is adapted from Melands and Willassen 2007. Decapod phylogeny is adapted from Scholtz and Richter 1995 and Schram 2001. Representative species are shown at each branch. Species denoted in purple are edible food crops.

We have annotated innate immunity genes and pathways from 69 Malacostraca transcriptome datasets from 55 species representing five Malacostraca orders: Amphipoda (7 species), Decapoda (18 species), Isopoda (27 species), Euphausiacea (2 species) and Mysida (1 species) (fig. 1; supplementary tables 1 and 2; Scholtz and Richter 1995; Schram 2001; Melands & Willassen 2007). We used sequence, motif and domain similarity based approaches to identified 7407 genes, representing 39 immune gene families in the Malacostraca (summarised in fig. 2). We annotate genes that encode pathogen recognition proteins, signalling components of key signal transduction pathways such as Toll, Imd and JAK-STAT, effector genes encoding proteins that perform immune protection such as antimicrobial peptides and members of the antiviral RNAi pathway. Within these key groups, we define malacostracan specific evolutionary events that suggest a previously unsuspected variation in immune gene content, and that functional genomic studies of immunity specifically with species in this group will be required for clear understanding of host defence in food crop species. A comparison across these data sets also allowed us to expand the annotation of previously discovered crustacean specific immune components, confirming their importance across the group. Finally, taking a conservative approach using orthology analyses and Pfam annotations in the sequenced amphipod genome of the crustacean ***P. hawaiensis*** (Kao et al, 2016), we describe four novel gene families with immune related protein domains conserved only within the Malacostraca. We show that these novel Malacostraca genes exhibit tissue-specific expression in the amphipod ***P. hawaiensis.*** Overall, our work provides a comprehensive picture of the Malacostraca innate immune system and a key staging point that will now facilitate important immunology research to underpin food crop aquaculture.

**Figure 2.**
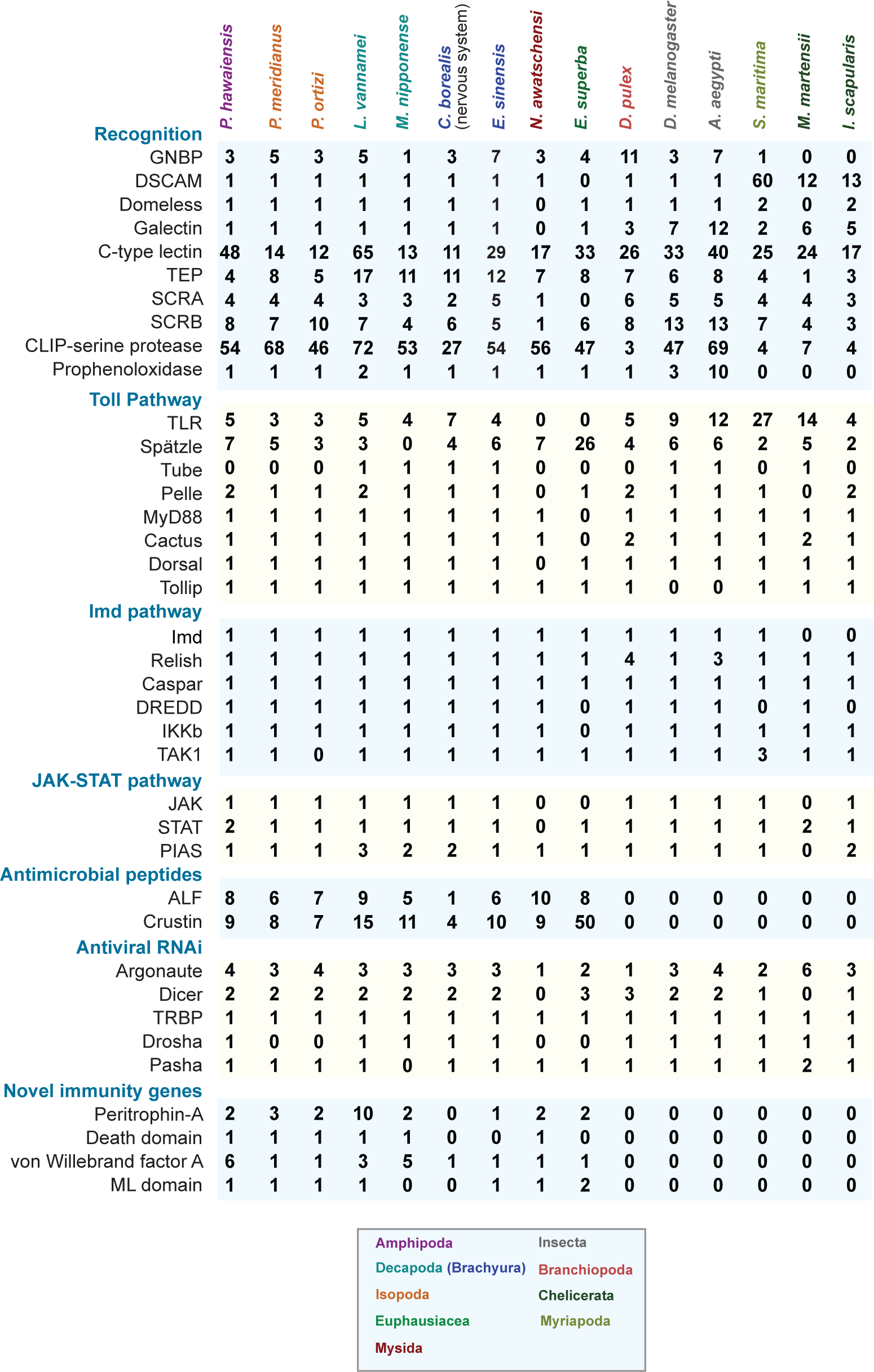
Summary of gene copy number in representative malacostracans and arthropods. Gene copy number for malacostracans are determined in this study. Gene copy number for arthropods were obtained from Waterhouse et al. 2007; McTaggart et al. 2009; Palmer and Jiggins, 2015 and Kao et al. 2016. A complete list of immunity genes identified in this study is presented in supplementary tables S3-S9 and supplementary files S1-S42.

## Results and Discussion

### Pattern recognition receptors in malacostracans are dynamically evolving and exhibit family-specific expansions

While vertebrates rely on adaptive immune systems and immunological memory mediated by secreted antibodies to ward off pathogens, many invertebrates, including arthropods use a pre-encoded set of proteins known as the pattern recognition receptors (PRRs) to recognise a broad spectrum of microbial ligands. Arthropods PRRs facilitate microbial killing through a range of direct and indirect mechanisms (De Gregorio et al., 2002; Hoffmann, 2003; Meister, 2004; Govind et al., 2004; Theopold et al., 2004; Brennan et al., 2004) upon the detection of non-self pathogen structures known as pathogen-associated molecular patterns (PAMPs) present on the surface of microbes (Janeway, 1989). Some examples of PAMPs include peptidoglycans (PGN) and lipotechoic acids (LTA) in Gram-positive bacteria, lipopolysaccharides (LPS) in Gram-negative bacteria and (β-glucans from fungal cell walls (Janeway, 1989; Beutler, 2004). We examined seven PRR families in Malacostraca, which included the Gram-negative binding proteins (GNBPs), Down syndrome cell adhesion molecules (DSCAMs), scavenger receptors (SRs), Domeless proteins (discussed in the signal transduction section), C-type lectins (CTLs), galectins and thioester-containing proteins (TEPs; fig. 2; fig. 3A). DSCAM, SRs and Domeless are transmembrane receptors (fig. 3A). GNBPs can either be associated with the cell membrane via a glycosylphosphatidylinositol anchor or function as soluble receptors (Ferrandon et al. 2007). We identified 202 GNBPs, 49 DSCAMs, 443 SRs, 47 Domeless proteins, 1005 CTLs, 47 galectins and 432 TEPs in malacostracans (supplementary table 3).

**Figure 3.**
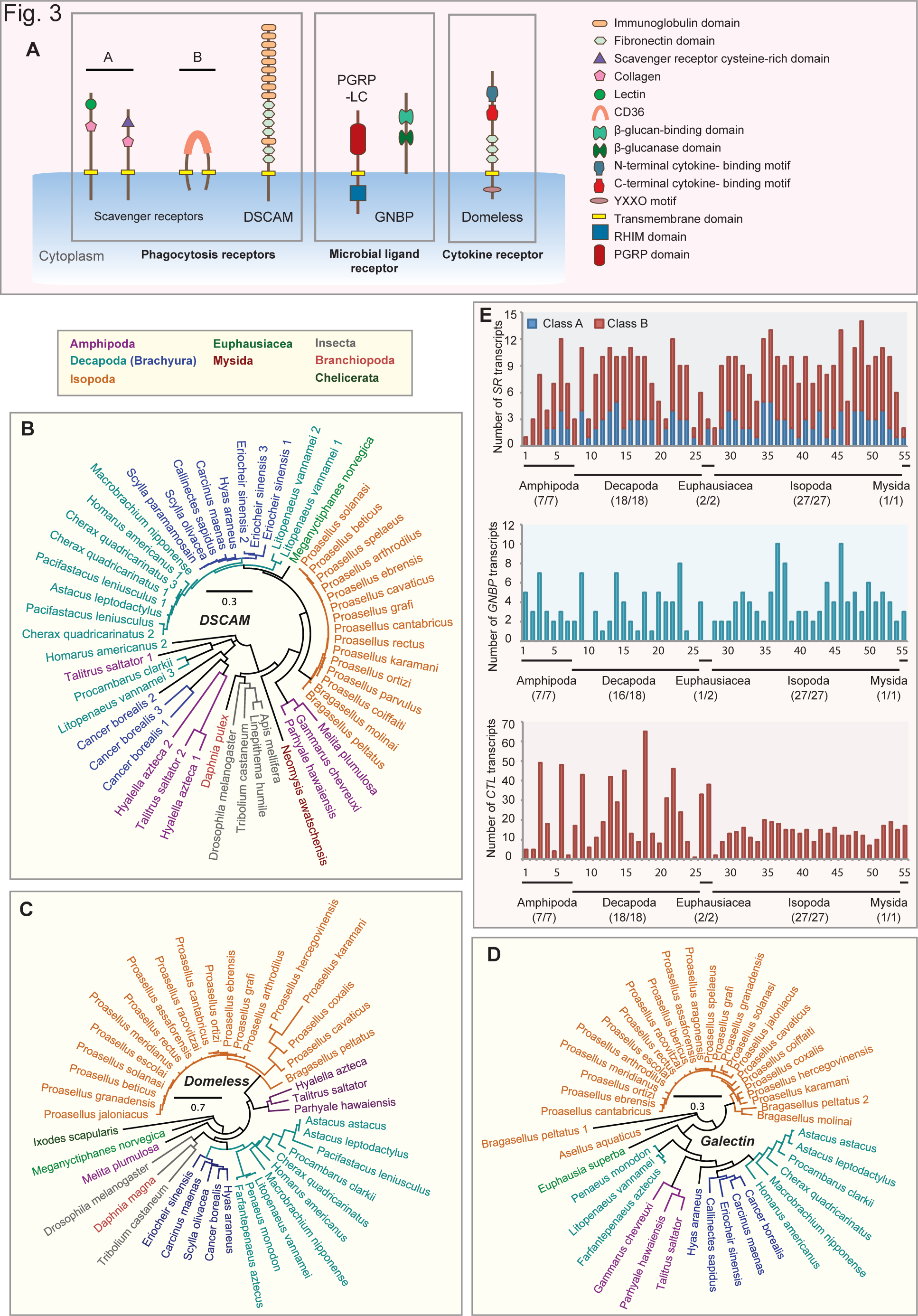
Innate immunity pattern recognition receptors (PRRs) in malacostracans. **(A)** Domain architecture of PRRs. The neuronal transmembrane protein of the immunoglobulin (Ig) superfamily DSCAM contains tandem arrays of Ig and fibronectin domains. DSCAM is shown to participate in pathogen recognition in mosquitoes and phagocytosis flies. Scavenger receptors are a diverse group of multidomain proteins. Two classes of the membrane-associated scavenger receptors are shown. Members of the class A scavenger receptors (SCRAs) subfamily contain the characteristic scavenger receptor cysteine-rich (SRCR) domain, C-type lectin and collagenous domains. Class B scavenger receptors (SCRBs) are characterised by two transmembrane domains and a CD36 domain. PGRP-LC contains the RHIM (receptor-interacting protein homotypic interaction motif) domain. GNBPs typically possess the β-glucan binding domain and the β- glucanase domain. Domeless is a cytokine receptor required for JAK-STAT signalling. Conserved protein domains of PRRs are shown in the figure inset. Phylogenetic trees of **(B)** DSCAM, **(C)** Domeless and **(D)** Galectin are constructed using the maximum-likelihood method from an amino acid multiple sequence alignment. Taxa labels are depicted as their respective colour codes. Bootstrap support values (n=1000) for all trees can be found in Supplementary figure 8. Scale bar represents substitution per site. **(E)** The graphs represent the repertoire of putative PRR transcripts of the following gene families: ***SCRAs and SCRBs, GNBPs*** and ***C-type lectins.*** The y-axes represent total number of genes identified in all 55 malacostracan species for each family. Each species is represented by a number on the X-axes and a complete list of species is available in Supplementary table 2. Black horizontal bars below each graph delimit the five orders of malacostracans and the numbers in parentheses (x/y) represent the following: x = number of species in which a particular gene family is found and y = total number of species in each order.

GNBPs are a group of proteins that share the carbohydrate-binding β-glucanase domain. Multiple naming conventions exist for this group as they are also called lipopolysaccharide and β-glucan binding proteins (LGBPs), β-1,3-glucan binding proteins (BGBP) or β-1,3-glucanase-related proteins (BGRP). Originally discovered in lepidopterans as proteins that can recognise β-1,3-glucans from fungal cell walls (Ochiai and Ashida 1988; Ochiai et al. 1992; Ochiai and Ashida 2000), others have shown that insect GNBPs can also recognise Gram-negative bacteria (Lee et al., 1996; Zhang et al., 2003; Pauchet et al., 2009; Genta et al., 2009; Bragatto et al., 2010). Functional studies on malacostracan GNBPs revealed that these proteins are expressed in hemocytes and hepatopancreas and expression is induced upon treatment with WSSV, Gram-negative and - positive bacteria (Roux et al., 2002; Sritunyalucksana et al., 2002; Cheng et al., 2005; Du et al., 2007; Lin et al., 2008; Liu et al., 2009). GNBPs have two active sites, identified in the p-glucanase domain of the silk moth ***Bombyx mori*** that are denoted as E188 and E193 (Zhang et al., 2003). Both glutamic acid residues are reported to be absent from most insect GNBPs, which implied that insect GNBPs lack catalytic activity (Zhang et al., 2003; Hughes, 2012). GNBPs have undergone significant evolutionary changes within Arthropoda because chelicerates lack GNBPs altogether (Palmer et al., 2015) while ***Drosophila melanogaster*** and ***D. pulex*** have 3 and 11 GNBP proteins respectively (fig. 2; supplementary table 3; McTaggart et al., 2009). We identified 202 GNBP genes from Malacostraca (supplementary table 3; fig. 3E). Of these, 150 have intact p-glucanase domains. Alignment of these β-glucanase domains from Malacostraca with the ***B. mori*** sequence revealed that 109 out of 150 malacostracan GNBPs possessed glutamic acid residues at both E188 and E193 positions (supplementary figure 1). From our analyses we show that a clear GNBP expansion has occurred within Malacostraca (fig. 3E). Many decapods have expanded GNBPs; for example ***Procambarus clarkii, Eriocheir sinensis*** and ***Astacus leptodactylus*** have 8, 7 and 7 GNBPs respectively (fig. 3E; supplementary table 3). GNBP expansion more broadly in Crustacea may compensate for the previously reported absence of peptidoglycan recognition proteins (PGRPs) in the crustacean lineage (McTaggart et al., 2009) since GNBPs can also recognise Gram-positive bacteria. While we do find some PGRPs for the first time in our analysis (see Imd signalling section), our data supports this idea. The 3 GNBPs in ***D. melanogaster*** are catalytically inactive (Palmer et al., 2015) and given that this species has 13 PGRPs, it is possible that in ***D. melanogaster***, PGRPs may compensate for inactive GNBPs. Thus in the malacostracan and insect lineages within Pancrustacea, these to key pathogen detection systems have undergone opposing evolutionary trajectories. This observation supports the suggestion that detailed functional genomic studies of immunity genes are required in a malacostracan species to properly make progress in food crop species immunology research.

A major component of arthropod immune systems that still requires further definition and is yet to be fully exploited is the DSCAM proteins, which undergoes startling levels of alternative splicing (AS) in the Pancrustacean clade. The canonical DSCAM domain arrangement consists of 9(Ig) - 4(fibronectin; Fn) - (Ig) -2(Fn) (Shapiro et al., 2007). Since other Ig-containing genes may confound the identification of bona fide ***DSCAM*** transcripts in malacostracans, we searched for genes/transcripts containing the Fn1-Fn2-Fn3-Fn4-Ig10-Fn5 motif set from known DSCAM protein sequences used as queries for BLAST. From this, we identified putative ***DSCAM*** transcripts in 49 out of 55 malacostracan species in our study (fig. 3B). We observed that DSCAMs in brachyurans (except for ***Cancer borealis)*** are monophyletic and are likely to be orthologous (fig. 3B). DSCAM AS in arthropods has evolved to allow versatile pathogen recognition, greatly increasing the reservoir of receptor diversity (Schmucker et al., 2000; Graveley et al., 2004; Dong et al., 2006; Brites et al., 2008; Yue et al., 2016; Kao et al., 2016), Although it is likely that most malacostracan ***DSCAMs*** have multiple splice forms, accurate characterisation and annotation of splice variants from transcriptome data alone is confounded by long arrays of highly similar Ig exons. Genome and genomic DNA based approaches will be needed to assess this with accuracy. Nonetheless is seems likely that the DSCAM remains a key PRR in malacostracans, and will need to be further studied in the context of infection as potential diagnostic marker and effector mechanism that might be exploited in aquaculture.

Scavenger receptors (SRs) are a subclass of structurally diverse membrane-bound PRRs, first described as proteins having the ability to bind to oxidised low-density lipoproteins (LDLs) (fig. 3A; Brown et al., 1979; Janeway, 1989; Krieger, 1997; Medzhitov et al., 2002; Areschoug et al., 2009; Canton et al., 2013). SRs can recognise a diverse range of cognate ligands and these include modified self-molecules (eg: oxidised LDLs) and non-self microbial structures such as LPS and LTA (Plüddemann et al., 2006; Plüddemann et al., 2011). We considered two classes of SRs in Malacostraca, namely the macrophage class A scavenger receptors (SCRAs) and the class B scavenger receptors (SCRBs; Fig. 3A). We annotated 129 SCRAs and 314 SCRBs in malacostracans (supplementary table 3; fig. 3E). Malacostracans SCRAs are characterised by multiple domains; the cysteine-rich (SRCR) domain, C-type lectin domain, lysyl oxidase (LOX) or collagen domain (fig. 3A; Hampton et al., 1991; Dunne et al., 1994; Peiser et al., 2002). Malacostracans SCRBs have the CD36 domain and two transmembrane domains (fig. 3A; Canton et al. 2013). To date, the only SR reported in crustaceans is a homolog of Croquemort, a SCRB family member in ***Marsupenaeus japonicus*** (Mekata et al. 2011). Humans and ***Caenorhabditis elegans*** only have three CD36-like proteins each (Febbraio et al., 2001; Stuart et al., 2008). SCRBs in malacostracans have however, undergone multiple gene duplications; the isopod ***Proasellus ortizi, the*** decapod ***C. borealis*** and amphipod ***P. hawaiensis*** have 10, 6 and 8 SCRBs respectively (fig. 2; supplementary table 3). Major SCRB gene expansion is likely to have occurred at the base of Mandibulata as ***S. maritima, D. melanogaster*** and ***D. pulex*** have 7, 13 and 8 homologs respectively while the chelicerate ***I. scapularis*** only has three (fig. 2). Clearly the role of SCRBs in mandibulate immunity needs further study as almost nothing is known about the significance of the SCRB expansion in arthropods. Perhaps not all SCRBs in malacostracans are involved in host defence because CD36-like proteins have been shown to participate in other physiological roles such as facilitating cellular uptake of carotenoids required for visual chromophore formation (Kiefer et al. 2002), scavenging of apoptotic cells (Franc et al. 1996) and lipoprotein homeostasis (Rigotti et al. 1995).

Lectins have been shown to be directly relevant to the immune system of crustaceans (Wang et al., 2013; Wongpanya et al. 2016). A C-type lectin in ***M. japonicus***, expressed primarily in intestinal tissues, is upregulated upon bacteria and WSSV infection and can bind LPS and PGN in a dose-dependent manner (Feng et al., 2016). Nonetheless little is known about how many of each of the different types of lectins are present in malacostraca. C-type lectins (CTLs) are a group of diverse proteins characterised by a carbohydrate-recognition domain, some of which are Ca^2^+^−^ dependent and they can bind sugar and non-sugar ligands (Drickamer and Taylor, 1993; Drickamer and Fadden, 2002), while galectins are another type of lectin proteins that can bind β-galactoside sugars and are involved in multiple cellular processes such as apoptosis, cell proliferation and immunity (Cooper et al., 1999). We identified over a thousand putative CTLs across Malacostraca with ***Litopenaeus vannamei*** having 65 different CTLs, in line with a general trend for decapods to have more CTLs than other malacostracan groups (fig. 2; fig. 3E; supplementary table 3). Although the copy number of CTLs varies greatly between malacostracan species (fig. 3E; supplementary table 3), it is clear that divergent evolution through multiple gene duplications has occurred within this lineage, particularly in some amphipod and decapod species (fig 3E). More broadly we find that this appears to be a feature in many other arthropod lineages; ***S. maritima, D. melanogaster, Aedes aegypti*** and ***D. pulex*** have 25, 33, 40 and 26 genes respectively (supplementary table 3). To date, only five CTLs in ***L. vannamei*** have been studied in the contexts of Gram-negative bacteria agglutination and WSSV infection (Ma et al. 2007; Sun et al. 2007; Zhang et al., 2009; Zhao et al. 2009; Wei et al. 2012). Future expression panel testing, particularly in decapods, will be required to ascertain whether CTLs may have distinct roles in recognising different pathogenic agents. Galectins in malacostracans are present as single-copy homologs except in two isopod species ***(Asellus aquaticus*** and ***Bragasellus peltatus*** that have two galectins each; supplementary table 3; fig. 3D). Our analysis revealed that insects, chelicerates, ***S. maritima*** and ***D. pulex*** have multiple copies of galectins (fig. 2) suggesting that with respect to galectins, malacostracans have evolved conservatively (fig. 3D).

The thioester-containing protein (TEP) superfamily includes the vertebrate complement system, the pan-protease inhibitor α2-macroglobulin (α2M), insect TEP-like proteins and macroglobulin complement related (MCR) proteins (Nonaka et al., 2004). TEPs have the unique propensity to form covalent bonds with pathogens through their canonical thioester (GCGEQ) motifs to promote endocytotic clearance or to neutralise pathogenic proteases (Blandin et al., 2004; Stroschein-Stevenson et al., 2006; Armstrong, 2010). Amongst arthropods, some TEPs lack the canonical thioester motif and they presumably lack the ability to form covalent bonds with pathogenic surfaces (Palmer et al., 2015). We identified a total of 432 TEPs in all 55 malacostracan species (supplementary table 3). Decapods in general have more TEPs than other malacostracan orders (supplementary figure 2B). ***P. clarkii*** has at least 25 different TEPs, the highest amongst the malacostracan datasets considered here (supplementary table 3). However, only a third (147/432) of malacostracans TEPs have the GCGEQ motif. Nonetheless, as reports have indicated that a TEP protein in ***D. melanogaster***, although lacking the thioester motif, could still bind to fungi and promote phagocytosis (Stroschein-Stevenson et al., 2006), this TEP diversity in malacostraca and particularly Decapoda may be immune related. We analysed phylogenetic relationships between TEP members in Malacostraca and observed that like TEPs in arthropods, they fall into three major categories: a2Ms, insect TEP-like proteins and MCRs (supplementary figure 2). The α2Ms in amphipods (except for 1 gene in ***Talitrus saltator)*** form a monophyletic group (supplementary figure 2A). The vertebrate C3 and C4 factors are also monophyletic, while C3 from the amphioxus ***Branchiostoma belcheri*** and C5 factors from mouse and human are paraphyletic (supplementary figure 2). Since we did not find any malacostracan TEPs clustering with the vertebrate complement factors and together with the observation that C3-like proteins are only found in chelicerates and myriapods (Palmer et al. 2015), we predict that C3-like proteins have been lost in the Pancrustacea.

The recognition of PAMPs by PRRs is the first line of defence against invading pathogens. In this study, we have annotated known PRR families in Malacostraca and established analogies to arthropod PRRs (fig. 2 and fig. 3). We show that malacostracans have a large repertoire of PRR proteins to efficiently cope with a broad range of pathogens. Several PRR families, CTLs, GNBPs and SCRBs, are expanded in malacostracans and this may, in part, contribute to enhanced plasticity when dealing with diverse microbial ligands. Our data will underpin comparative approaches as to how PRR activation in aquaculture affects outcomes in different conditions. Together our analyses indicate that PRRs are evolving rapidly within this lineage, reflecting the diverse selection pressure from pathogens encountered by different malacostracan groups.

### Prophenoloxidases are invented at the base of Pancrustacea

The prophenoloxidase-activating system (proPO) is another non-self pathogen recogonition mechanism implicated in arthropod immunity (Söderhäll et al., 1994; Söderhäll and Cerenius 1998; Sritunyalucksana & Söderhäll, 2000). Upon the recognition of LPS, PGNs or β-glucans by GNBPs, a serine protease cascade ensues, which results in the proteolytic cleavage of proPO into active phenoloxidase (PO). PO then catalyses melanin formation (Söderhäll and Cerenius 1998) and this creates a physical barrier to inhibit further pathogen growth and movement (Cerenius and Söderhäll 2004). Because PO plays functional roles in the melanisation pathway and wound healing (Áspan et al., 1991; Söderhäll et al., 1998; Ashida et al., 1998), the emergence of PO is associated with the evolution of humoral immunity in arthropods. POs are thought to be members of the hemocyanin superfamily; a family that is exclusively found in arthropods (Burmester, 2002). Due to shared sequence similarities, it was proposed that hemocyanins could be converted to proPOs upon chemical treatments (Decker et al. 2001; García-Carreño et al. 2008). Chelicerates (scorpions and spiders) and the myriapod ***S. maritima*** lack ***sensu stricto*** proPOs (fig. 2; Burmester, 2002; Jaenicke et al., 2004; Decker et al., 2007; McTaggart et al., 2009; Palmer et al., 2015) and so perhaps they would need to rely on activated hemocyanins for melanin synthesis. To date, most crustacean proPOs were identified from decapods (Hernández-López et al. 1996; Lee et al. 2004; Ko et al. 2007; Cerenius et al. 2008; Masuda et al. 2012; Amparyup et al. 2013; Masuda et al. 2014). We found only two malacostracan proPOs from non-decapod species in GenBank, ***Nebalia kensleyi*** (Leptostraca; ACV33307.1), and ***Oratosquilla oratoria*** (Stomatopoda; ADR50356.1; supplementary figure 3A), indicating that proPO exists beyond decapod species. No other reports exist for proPOs in amphipods (except for ***P. hawaiensis;*** Kao et al. 2016), isopods, krills and mysid crustaceans. Some have reported that amphipods and isopods lack proPO (Pless et al., 2003; Hagner-Holler et al., 2005; Terwilliger et al., 2006; Jaenicke et al., 2009; King et al., 2010). Failure to identify proPOs from isopods by an independent study could be due to the use of a limited EST dataset (King et al., 2010). In contrast to the previous studies, we were able to identify proPOs from all five malacostracan orders (supplementary figure 3A). Since proPOs and hemocyanins have similar sequences, we confirmed that these are ***bona fide*** proPOs through reciprocal BLASTs and phylogenetic analysis (supplementary figure 3A). Considering that proPOs are present in insects, ***D. pulex*** and malacostracans but not in myriapod and chelicerate lineages (although related proteins with predicted tyrosinase activity were identified; Palmer et al., 2015), it is likely that this non-oxygen binding derivative of hemocyanin was invented at the base of Pancrustacea. ProPOs may have evolved distinct roles in immunity since we were still able to identify many other hemocyanin genes in malacostracans (supplementary figure 3B). Parallels have been drawn between the initiation of serine protease cascades and the conversion of proPOs into catalytically active POs after exposure to PAMPs (Kanost et al. 2004; Tang et al. 2006). POs must be tightly regulated by serine proteases since PO activation generates highly reactive toxic quinone intermediates (Smith & DeLotto, 1992; Jiang et al., 1998; Cerenius and Söderhäll 2004) and CLIP-domain serine proteases are implicated in this process (Jiang & Kanost, 2000). CLIP-domain serine proteases are expanded in Diptera insects ***(D. melanogaster*** has 47 genes) but not in ***D. pulex, S. maritima*** and chelicerates (fig. 2; Waterhouse et al., 2007; Palmer et al. 2015). We made similar observations on the expansion of CLIP serine proteases in Malacostraca (supplementary figure 3C). We annotated over 2163 CLIP serine proteases. The highest numbers across five malacostracan orders are: the decapod ***L. vannamei*** (72 genes), the amphipod ***P. hawaiensis*** (54 genes), the isopod ***Proasellus meridianus*** (68 genes), the krill ***Meganyctiphanes norvegica*** (57 genes) and the mysid crustacean ***Neomysis awatschensis*** (56 genes; supplementary table 3). The expansion of CLIP serine proteases in malacostracans may signify a need for highly regulated PO activation and this correlates with our novel findings of proPO presence across the broader Malacostraca.

### Toll and JAK-STAT pathways are conserved in Malacostraca while several key components of the Imd pathway are lost

Signal transduction pathways link recognition of PAMPs by PRRs with transcriptional activation. Three well-studied pathways are the Toll, Immune Deficiency (Imd) and Janus Kinase (JAK)-signal transducer and activators of transcription (STAT) pathways. Components of the Toll pathway in malacostracans include a chain of interacting proteins: the Toll-like receptors (TLRs; Akira et al., 2006; Pasare et al., 2005; Leulier et al., 2008), Spätzle (Lemaitre et al., 1996; Rutschmann et al., 2002; Shi et al., 2009; Wang et al., 2012), myeloid differentiation factor 88 (MyD88; Tauszig-Delamasure et al., 2002), Tube, Pelle, Dorsal, Cactus and the Toll-interacting protein (TOLLIP; Fig. 4A). Our investigation of the malacostracan Toll pathway members suggests that this pathway is broadly conserved. Malacostracan TLRs appear to have undergone divergent evolution through multiple gene duplications and from our phylogenetic analysis, we saw that paralogs exhibited marked sequence divergence (supplementary figure 4). Like in dipterans ***(D. melanogaster*** and ***A. aegypti*** have 6 genes each), we discovered multiple-copies of the gene encoding Spätzle, the cytokine partner of Toll, in malacostracans. In particular we observed that ***Euphausia superba*** has at least Spätzle encoding genes, while we found numbers more in line with insects in ***E. sinensis*** (6 genes), ***A. leptodactylus*** (6 genes), ***P. hawaiensis*** (7 genes), ***M. norvegica*** (5 genes), ***N. awatschensis*** (7 genes) and ***Proasellus sonalasi*** (7 genes; fig. 4B). We conclude that the unique expansion in ***E. superba*** might be a unique curiosity of this species. We identified intact MyD88-Tube-Pelle complexes in only 13 out of 55 malacostracan species (fig. 4C). As in insects (Christophides et al., 2002; Waterhouse et al., 2007), MyD88 and Tube each exist as single-copy genes in Malacostraca (supplementary table 4). We found Tube transcripts from 17 species representing only the Decapoda and Isopoda orders. Only one report of a crustacean Tube homolog has been previously shown (Li et al., 2014). Although dipterans have one copy of Pelle, we observed duplications of Pelle in some malacostracan species in the Decapoda ***(Homarus americanus*** and ***L. vannamei)***, Amphipoda ***(Echinogammarus veneris, P. hawaiensis*** and ***T. saltator)*** and Isopoda ***(Proasellus beticus, P. coiffaiti, P. coxalis, P. grafi, P. hercegovinensis*** and ***P. rectus)*** (fig. 4C; supplementary table 4). As two copies of Pelle were found in ***D. pulex*** and the deer tick ***Ixodes scapularis*** (Palmer et al., 2015), it is possible that duplication of Pelle may have occurred at the base of arthropod lineages, with subsequent loss of one copy in insects. We identified single-copy homologs of Dorsal and Cactus in malacostracans (fig. 4D, F and G). The Toll-interacting protein (TOLLIP) is a negative regulator of NF-κB in mammals (Zhang et al., 2002). Not much is known about the function of TOLLIP in invertebrates and to date, only one TOLLIP in crustaceans has been described (Wang et al., 2013). We identified single-copy homologs of TOLLIP across five Malacostraca orders, ***D. pulex***, the myriapod ***S. maritima*** and chelicerates ***(Mesobuthus martensii*** and ***Ixodes scapularis)*** (supplementary table 4, Fig. 4E). Amongst dipterans, we identified single-copy TOLLIP homologs in ***Anopheles gambiae*** but neither in ***D. melanogaster*** nor ***A. aegypti***, although it is present in other insects like bees and ants (fig. 4E; supplementary table 4). Malacostracan TOLLIPs share similarities to mammalian TOLLIP proteins having both the protein kinase C conserved region 2 (C2) and the C-terminal coupling of ubiquitin to endoplasmic reticulum degradation (CUE) domain (Burns et al., 2000; Zhang et al., 2002).

**Figure 4.**
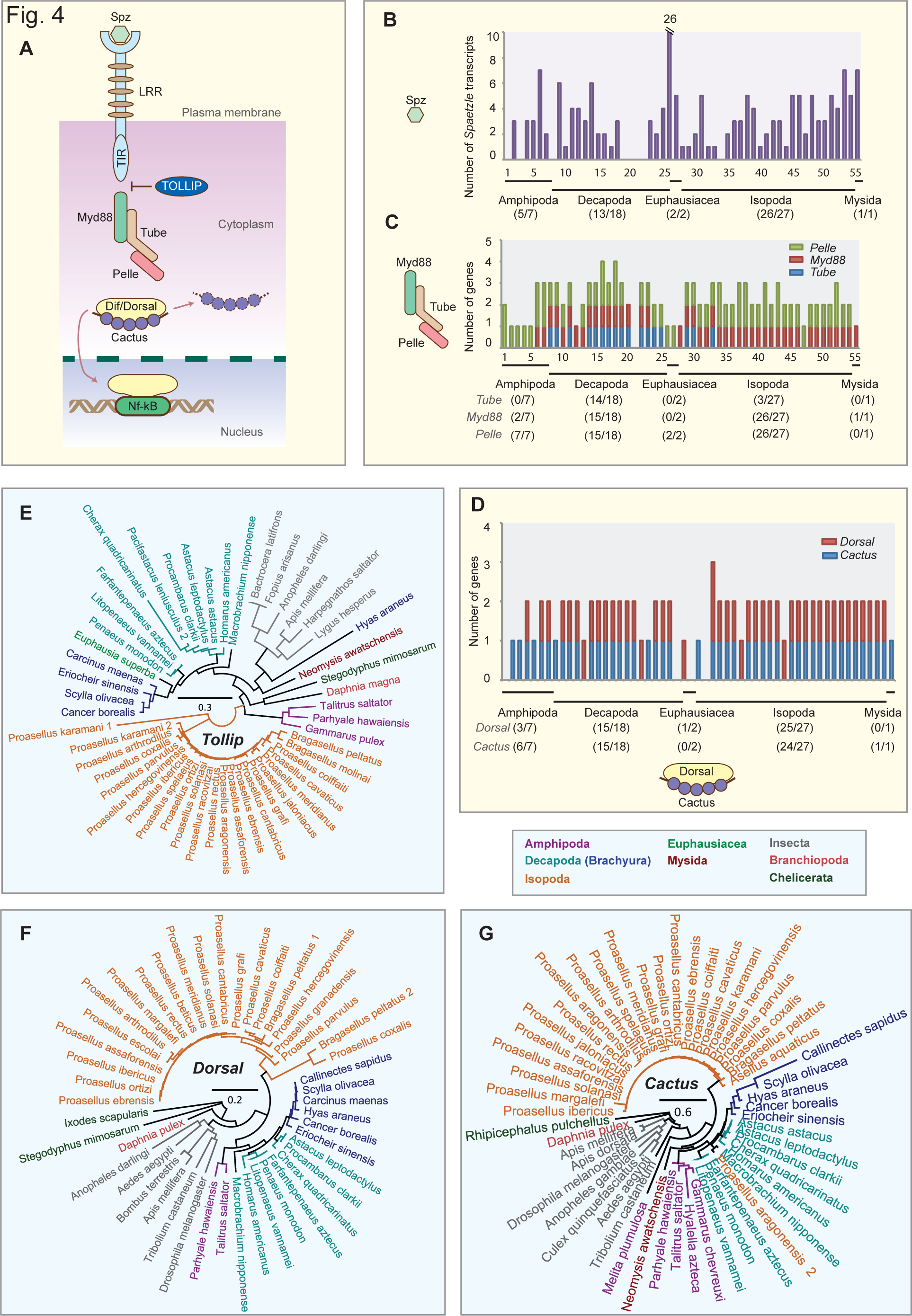
Toll pathway members in malacostracans. **(A)** Soluble PRRs such as the GNBP1 and GNBP3 are involved in the recognition of non-self, e.g. peptidoglycans and β-glucans, which triggers proteolytic through the activation of CLIP-domain serine proteases. The cytokine Spaetzle is cleaved by the Spaetzle processing enzyme and this activates the Toll receptor. Signalling through Toll acts via the Toll-induced signalling complex (TISC), comprising of three proteins containing death-domains: Tube, myeloid differentiation primary-response gene 88 (MyD88) and Pelle. TICS signal is transduced to Cactus (a homologue of the mammalian inhibitor of NF-κB). Cactus is phosphorylated, polyubiquitylated and degraded and the dorsal-related immunity factor (DIF) is translocated to the nucleus. DIF binds to NF-κB response elements to induce gene expression. The graphs represent the total number of **(B)** ***Spätzle***, **(C)** ***MyD88, Tube and Pelle*** and **(D)** ***Dorsal*** and ***Cactus*** transcripts in malacostracans. The y-axes represent total number of genes identified in all 55 malacostracan species for each family. Each species is represented by a number on the X-axes and a complete list of species is available in Supplementary table 2. Black horizontal bars below each graph delimit the five orders of malacostracans and the numbers in parentheses (x/y) represent the following: x = number of species in which a particular gene family is found and y = total number of species in each order. Phylogenetic trees of **(E)** Toll-interacting protein (TOLLIP), **(F)** Dorsal and **(G)** Cactus are constructed using the maximum-likelihood method from an amino acid multiple sequence alignment. Taxa labels are depicted as their respective colour codes. Bootstrap support values (n=1000) for all trees can be found in supplementary figure 8. Scale bar represents substitution per site.

Most components of the Imd pathway are present in malacostracans except for three gene families (fig. 5A). Imd is conserved amongst insects, myriapods and ***D. pulex***, but not in chelicerates (Bechsgaard et al., 2016; Palmer et al., 2015). Imd exists as a single gene within malacostracans across all five orders (Fig. 5B). Imd is preferentially activated by the inner PGN layer of Gram-negative bacteria through the binding of PGRP-LC to PGN (Gottar et al., 2002; Kaneko et al., 2006). To our knowledge, no PGRP homologs have been previously reported in crustaceans including ***D. pulex*** (McTaggart et al., 2009) and ***P. hawaiensis*** (Kao et al., 2016). Although we failed to identify PGRPs in most malacostracans, we found four putative PGRP genes from ***T. saltator*** (Amphipoda), ***Proasellus karamani*** (Isopoda) and ***H. americanus*** (Decapoda; supplementary figure 5A). This could indicate a complex pattern of PGRP loss amongst crustacean taxa, that PGRPs are present but not represented in available malacostracan transcriptome and/or that the PGRP sequences we have found have evolved convergently. Sequence analysis revealed that these malacostracans PGRPs possess the amidase domain and share striking sequence similarities to ***D. melanogaster*** PGRP-SC1, SC2 and SB2 (supplementary figure 5D). Within this domain, five amino acid residues (H-Y-H-T-C; marked in supplementary figure 5D) have been shown to be critical for PGRP enzymatic activity (Cheng et al., 1994; Mellroth et al., 2003). These residues are present in the malacostracans PGRPs annotated here, indicating that they have the potential to be enzymatically active. These data suggest that PGRPs are present in Crustacean taxa but perhaps have greatly reduced representation. Future genome sequenced based analyses will be required to clarify this. A negative regulator of Imd signalling is the Caspar protein, a homolog of the mammalian Fas-associating factor 1 (Kim et al. 2006). We identified single homologs of ***Caspar*** across all five malacostracan orders (Fig. 5C) and in other arthropods ***(D. pulex***, dipterans, chelicerates and myriapod) indicating that it is conserved in Arthropoda (supplementary table 5). Concerning other Imd pathway components, we identified single-copy homologs of Relish, death-related ced-3/Nedd2-like protein (DREDD),β IkB kinase β(IKKβ) and MAPKKK transforming growth factor -β (TGFβ)-activated kinase 1 (TAK1) in malacostracans indicating that these components of the Imd pathway have remained intact (fig. 5, supplementary figure 5 and supplementary table 5). However, we failed to identify clear homologs of IkB kinase y (IKKy), FAS-associated death domain (FADD) and TAK1-binding protein (TAB2) in malacostracan transcriptomes, which could be due to sequence divergence or the replacement of these components with other functionally related proteins. Independently, components of the Imd pathway have been reported in the decapod ***Carcinus maenas*** and the authors also did not find clear homologs for IKKγ or FADD (Verbruggen et al., 2015). Not much is known about the Imd pathway in crustaceans. To date, only two homologs, Imd and Relish have been subjected to functional studies in shrimps (Wang et al., 2009; Li et al., 2009; Huang et al., 2009; Wang et al., 2012; Li et al., 2013). Overall, the Imd pathway appears to be reduced in malacostracans. Whether this is a result of actual gene loss, sequence divergence or the utilization of alternative proteins for Imd signalling is presently unknown and warrants further investigation.

**Figure 5.**
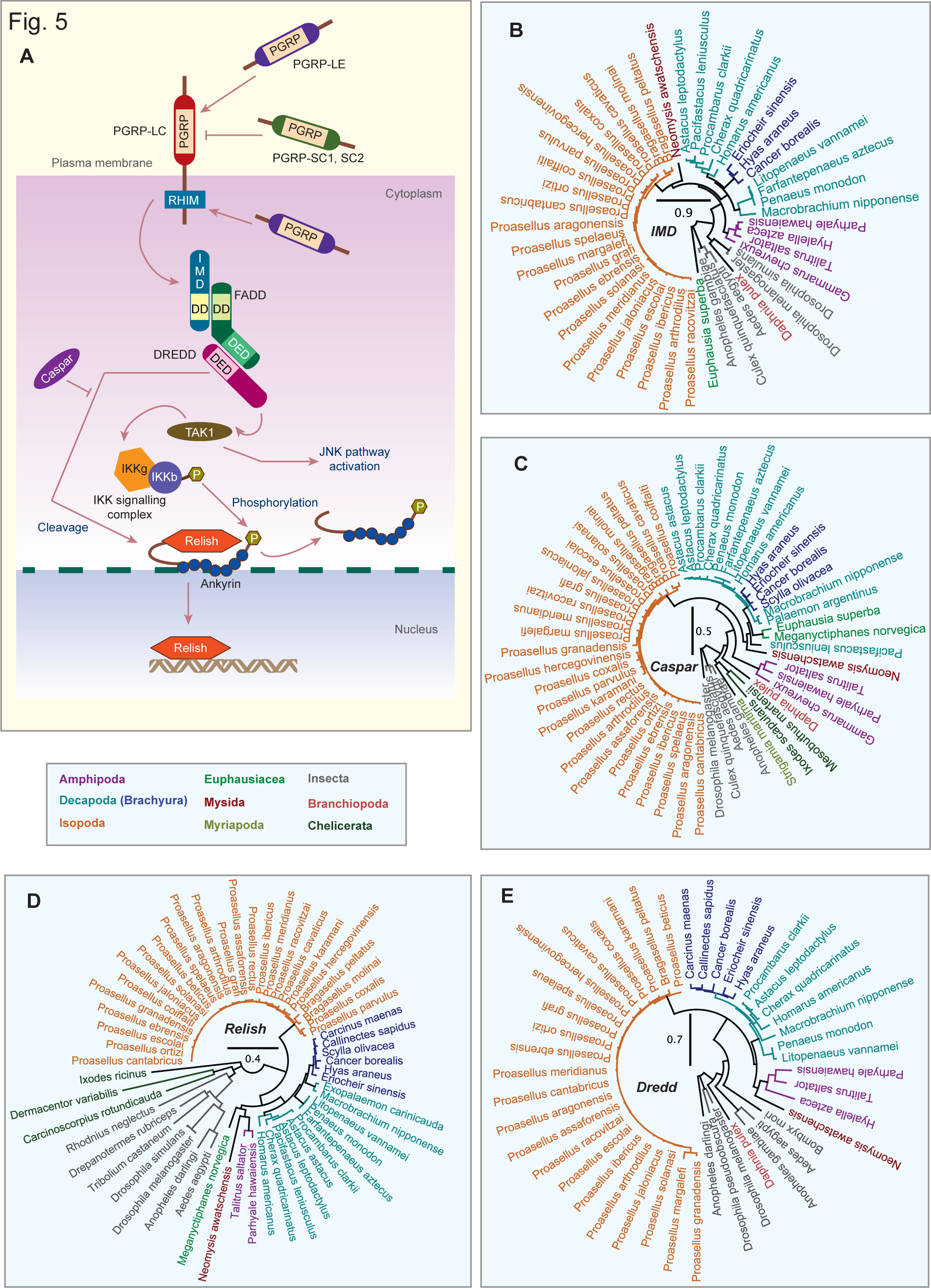
Immune deficiency (IMD) pathway members in malacostracans. **(A)** PGRPs recognises Gram-negative bacteria and activate the IMD pathway through the RHIM motifs. Although the IMD pathway is typically activated by PGRPs in ***Drosophila melanogaster***, PGRPs are not necessary for IMD signalling and it was posited that an unknown protein is present upstream of the IMD signalling cascade. Like DIF from the Toll pathway, in the IMD pathway, differential activation of another NF-κB transcription factor, Relish, occurs. Relish is phosphorylated through the activation of IκB kinase (IKK) complexes and transforming growth factor-β-activated kinase 1 (TAK1). The caspase-8 homolog death-related ced-3/Nedd2-like protein (DREDD) and FAS-associated death domain (FADD) proteins are required for IKK and TAK1 activation and Relish is cleaved through DREDD. Caspar, a homologue of mammalian Fas-associating factor 1 that is essential for antifungal immunity, negatively regulates the IMD-mediated immune response by preventing nuclear translocation of Relish. Caspar also suppresses the IMD pathway through targeting Dredd-dependent cleavage of Relish. Phylogenetic trees of **(B)** IMD, **(C)** Caspar, **(D)** Relish and **(E)** DREDD are constructed using the maximum-likelihood method from an amino acid multiple sequence alignment. Taxa labels are depicted as their respective colour codes. Bootstrap support values (n=1000) for all trees can be found in supplementary figure 8. Scale bar represents substitution per site.

Core components of JAK-STAT include the cytokine transmembrane receptor Domeless, JAK (Hopscotch in ***D. melanogaster)*** and STAT proteins (fig. 6A; Shuai et al., 2003; Arbouzova et al., 2006; Li, 2008; Morin-Poulard et al., 2013). Mammals have four JAK proteins and seven STATs (Levy et al., 2002) while most arthropods, except chelicerates, only have single-copy JAK and STAT homologs (Waterhouse et al., 2007; Palmer et al., 2015). JAK and Domeless proteins in malacostracans exist as single homoogs (fig. 3C; fig. 6B, supplementary table 6). Like Domeless, STAT in most malacostracans exists as single homologs, except in two amphipod species; ***Hyalella azteca*** (2 genes) and ***T. saltator*** (3 genes; fig. 6C, Supplementary table 6). Negative regulators of JAK-STAT include the suppressor of cytokine signalling (SOCS) and protein inhibitors of activated STAT (PIAS) (Yoshimura et al., 2007; Grönholm et al., 2010). SOCS and PIAS are also well conserved in malacostracans (supplementary table 6; fig. 6D and supplementary figure 6). Mammals have eight SOCS proteins while ***D. melanogaster*** only has three (Ilangumaran et al., 2004; Morin-Poulard et al., 2013). Copy number of SOCS varies between malacostracans; some of the highest numbers are in ***L. vannamei, P. hawaiensis*** and ***M. nipponense*** where they have 6, 6 and 5 genes respectively (supplementary table 6). Phylogenetic analysis of malacostracan SOCS proteins revealed that they clustered in seven major groups (supplementary figure 6). Few studies on crustacean SOCSs are available (De Zoysa et al., 2009; Zhang et al., 2010), and whether the entire malacostracan SOCS repertoire have roles in immunity is yet unknown.

**Figure 6.**
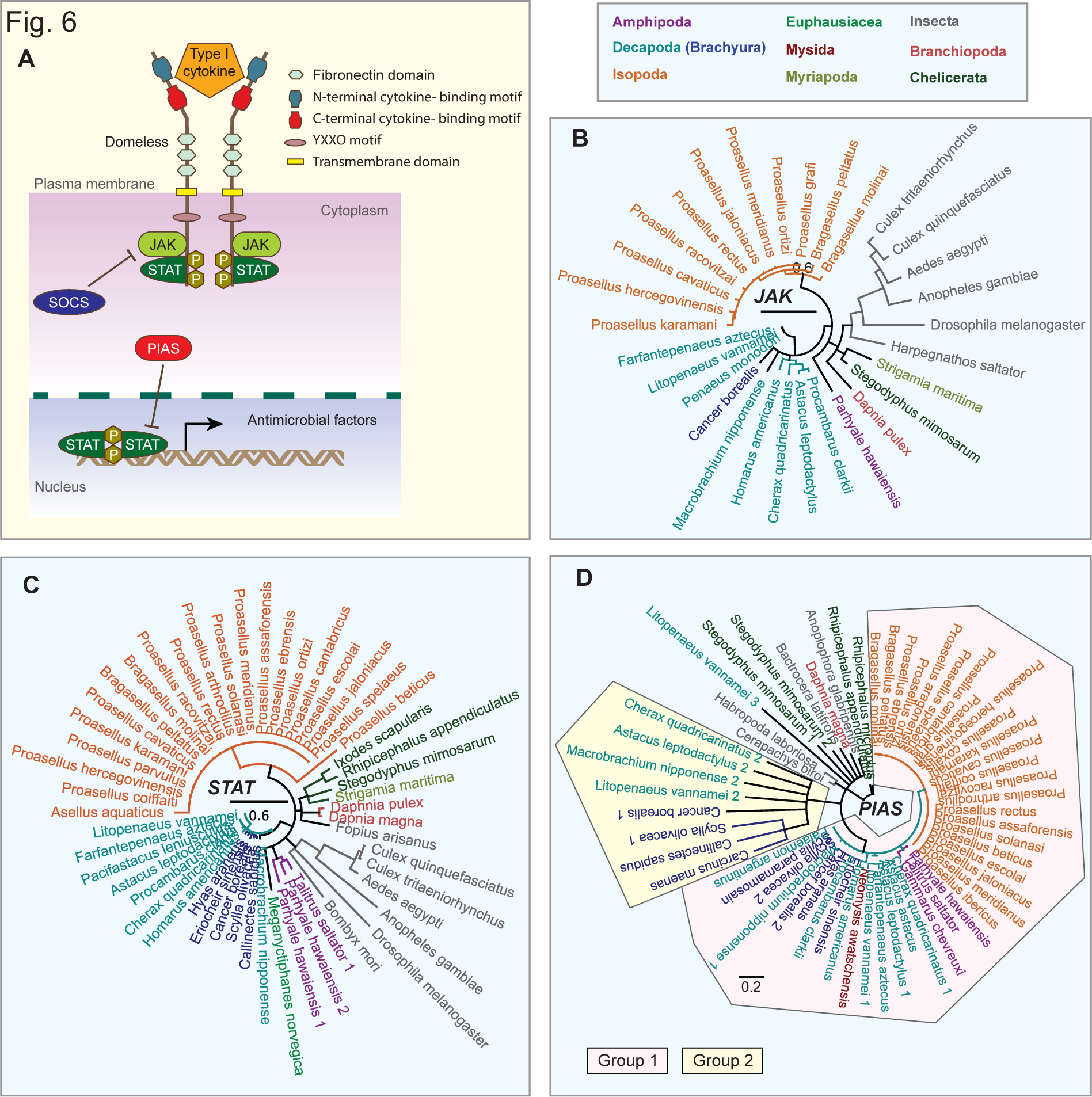
JAK-STAT pathway members in malacostracans. **(A)** Activation of the JAK-STAT signalling occurs through the binding of ligands such as cytokines to the cytokine receptor Domeless. Conserved protein domains of Domeless are shown in the figure inset. This binding activates the phosphorylation of the Janus kinase (JAK) proteins, which creates docking sites for Signal Transducer and Activator of Transcription (STAT) proteins through their Src Homology 2 (SH2) domains. STATs are phosphorylated by JAKs and activated STATs dimerise and are translocated to the nucleus to induce transcription. JAK-STAT transduction is controlled by suppressors of cytokine signalling (SOCS) and protein inhibitors of activated STAT (PIAS). SOCS proteins inhibit STATs phosphorylation via two mechanisms; 1) by competing with STATs for phosphotyrosine binding sites on cytokine receptors and (2) by binding to JAKs and preventing the recruitment of STATs onto the Domeless receptor. PIAS, also known as the E3 SUMO-protein ligase PIAS, is a transcriptional co-regulator that has the ability to inhibit STAT function. Phylogenetic trees of **(B)** JAK, **(C)** STAT and **(D)** PIAS are constructed using the maximum-likelihood method from an amino acid multiple sequence alignment. Two groups of PIAS proteins have been identified in malacostracans. Taxa labels are depicted as their respective colour codes. Bootstrap support values (n=1000) for all trees can be found in supplementary figure 8. Scale bar represents substitution per site.

Overall we have shown that three signal transduction pathways, Toll, Imd and JAK-STAT have remained largely conserved in malacostracans. Nonetheless, several components of these pathways exhibit lineage specific diversification, for example, the loss of three Imd pathway modules (IKKy, FADD and TAB2) and the divergent evolution of core TLRs and Spätzle components of the Toll pathway.

### Anti-lipopolysaccharide factors and crustins are malacostracan-specific antimicrobial peptides

Signal transduction culminates in the activation of immune effector molecules to neutralise pathogenic agents. Antimicrobial peptides (AMPs) are rapidly evolving, highly specific effector proteins that are potent agents against a broad range of microbes (Yount et al., 2006; Guani-Guerra et al., 2010). ***D. melanogaster*** has seven AMP families, but only three of these, attacins, cecropins and defensins, are shared with other dipterans (Waterhouse et al., 2007). To date, fifteen AMP families have been reported in crustaceans, fourteen of these are from decapods and many are lineage-specific (Gueguen et al., 2006; Rosa et al., 2010). We considered two of these AMP families, anti-lipopolysaccharide factors (ALFs) and crustins, and show that they are actually well conserved in malacostracans beyond just the Decapoda (fig. 7 and 8). While ALFs have only been reported in decapods (Zhao et al., 2008; Rosa et al., 2010) crustins have been reported once in a non-decapod malacostracan species, the amphipod ***Gammarus pulex*** (Smith et al., 2008). The branchiopod ***D. pulex*** lack ALFs and crustins or anything with sequence similarity, and we did not identify clear homologs in other arthropods, indicating that both gene families are specific to Malacostraca (supplementary table 7; fig. 7A). We identified a total of 337 ALFs from malacostracans form a wide range of tissue samples (supplementary table 7). The decapod ***Hyas araneus*** has the highest number of ALFs (20 genes; supplementary table 7; fig. 7B). Using homology modelling, we find that ALFs in malacostracans share high structural similarities, consisting of three β-sheets and three α-helices (fig. 7A'). Alignment analysis of malacostracan ALFs revealed that they contained two conserved cysteine residues predicted to form a disulfide bridge (Gross et al., 2001; Supungul et al., 2002; Rosa et al., 2010; fig. 7C). Between the cysteine residues, a region containing positively charged amino acids is defined as the LPS-binding domain (Zhao et al., 2008). This domain is present in all malacostracan ALFs, which suggests a conservation of LPS binding across this whole gene family (fig. 7C).

**Figure 7.**
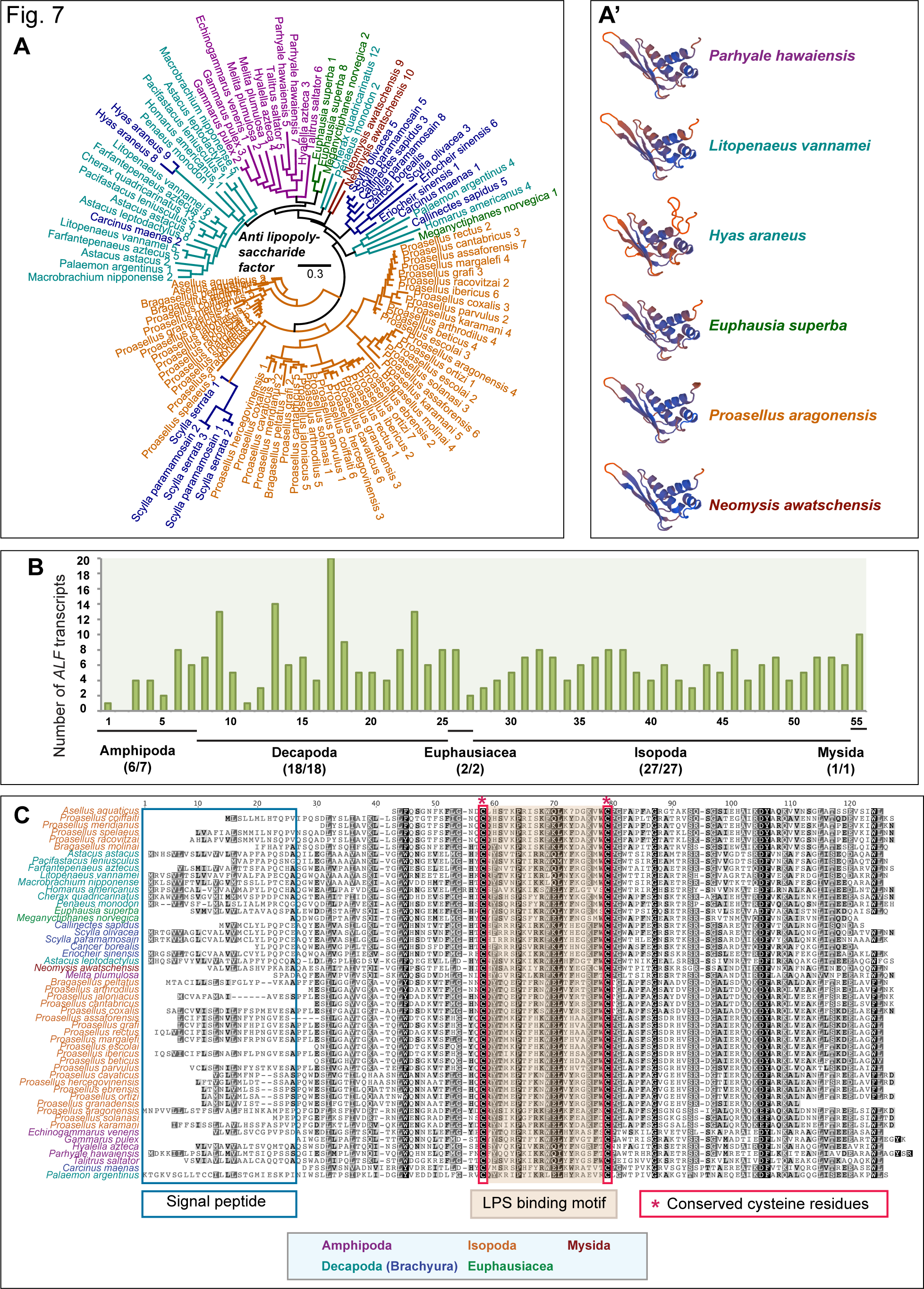
Anti-lipopolysaccharide factors (ALFs) in Malacostraca. **(A)** Phylogenetic tree of ALFs constructed using the maximum-likelihood method from an amino acid multiple sequence alignment. **(A')** Homology models of ALFs constructed with SWISS-MODEL revealed highly conserved predictions of the structural fold of these proteins. ALFs have conserved α-helical and β-strand structures. **(B)** Graph of putative ***ALF*** transcripts. The y-axes represent total number of genes identified in all 55 malacostracan species for each family. Each species is represented by a number on the X-axes and a complete list of species is available in Supplementary table 2. Black horizontal bars below each graph delimit the five orders of malacostracans and the numbers in parentheses (x/y) represent the following: x = number of species in which a particular gene family is found and y = total number of species in each order. Taxa labels are depicted as their respective colour codes. Bootstrap support values (n=1000) for all trees can be found in supplementary figure 8. Scale bar represents substitution per site. **(C)** Multiple sequence alignment of ALFs showing the putative signal peptides and LPS binding motifs characterised by two conserved cysteine residues are marked in red boxes.

Crustin is a cysteine-rich AMP containing a whey acidic protein (WAP) domain and was first discovered in the decapod ***Carcinus maenas*** to have a role in defence against Gram-positive bacteria (Relf et al., 1999). Crustins are abundant in malacostracans and we identified 513 putative genes with ***E. superba*** encoding at least 50 crustins (fig. 8A and B; supplementary table 7). In comparison with other WAP domain proteins, crustins are characterised by an additional crustin domain consisting of 12 conserved cysteine residues, in which a single WAP domain is present and we note that this the case in all malacostracans crustins (fig. 8C). Future studies can now address the biological roles of these AMPs and questions as to whether these AMPs are differentially regulated by specific microbial ligands, whether they are broad spectrum or selective and whether they are active in specific developmental stages. Our analyses show that the immune effector phase in malacostracans has undergone substantial lineage specific evolution, expansion and sequence diversification of AMPs, reflecting their modes of action to guard against a broad range of pathogens found in their natural habitats. We also note conservation of AMPs across Malacostraca, meaning that non-food crop species that can be easily studied in the lab will be potential model systems for this aspect of malacostracan immunity.

**Figure 8.**
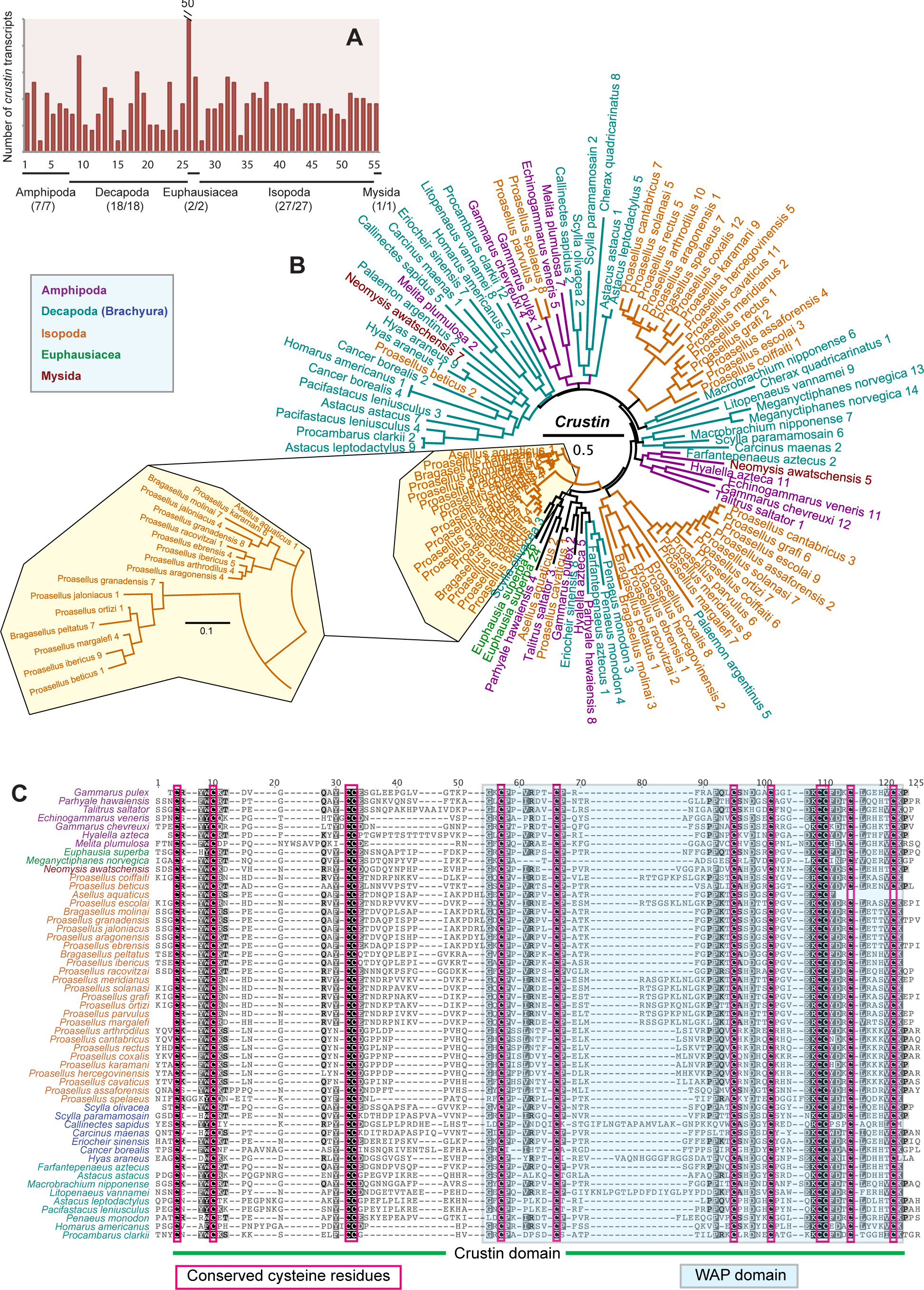
Crustin antimicrobial peptides in malacostracans. **(A)** Graph of putative ***crustin*** transcripts. The y-axes represent total number of genes identified in all 55 malacostracan species for each family. Each species is represented by a number on the X-axes and a complete list of species is available in Supplementary table 2. Black horizontal bars below each graph delimit the five orders of malacostracans and the numbers in parentheses (x/y) represent the following: x = number of species in which a particular gene family is found and y = total number of species in each order. **(B)** Phylogenetic tree of crustins is constructed using the maximum-likelihood method from an amino acid multiple sequence alignment. Taxa labels are depicted as their respective colour codes. Bootstrap support values (n=1000) for all trees can be found in supplementary figure 8. Scale bar represents substitution per site. **(C)** Multiple sequence alignment of crustins showing the crustin domain and the WAP domain within it. The WAP domain is characterised by 8 conserved cysteine residues marked in red boxes. The crustin domain, which includes the WAP domain contains four additional cysteine residues marked in red boxes.

### Malacostracans have a canonical RNAi-based antiviral immune system

RNA interference (RNAi) is a conserved antiviral mechanism in many systems (Lecellier et al., 2004; Li et al., 2005; Wang et al., 2006; Zambon et al., 2006; Cullen, 2006; Liu et al., 2009; Labreuche et al., 2012). RNAi-mediated gene silencing is now employed as a method to prevent viral disease progression in shrimps through the targeting of viral genes in order to inhibit replication (Robalino et al., 2005; Yodmuang et al., 2006; Kim et al., 2007; Xu et al., 2007; Tirasophon et al., 2007). No direct mechanistic evidence exists regarding the involvement of the RNAi pathway components in crustacean innate immunity. Despite this, there have been increasing efforts to identify RNAi pathway members in penaeid shrimps because of the potential applicability of RNAi-derived technologies in circumventing viral diseases (Robalino et al. 2005; Westenberg et al. 2005; Ding et al. 2007; Robalino et al. 2007; Wu et al. 2007; Xu et al. 2007; Liu et al. 2009; Labreuche et al. 2010; Bartholomay et al. 2012; Labreuche et al. 2013). We annotated core RNAi components in malacostracans, which include the RNAi-induced silencing complex (RISC) made up of three proteins: Dicer, the trans-activating response (TAR) RNA-binding protein (TRBP) and Argonaute-2 (fig. 9A). We found single-copy homologs of TRBPs across all five malacostracan orders and they share the conserved dsRNA-binding domain (fig. 9D; supplementary table 8). We identified Dicer proteins in amphipods, isopods, decapods and krills, but not in the mysid crustacean ***N. awatschensis*** (supplementary table 8; fig. 9B). Dicer-1 and Dicer-2 have distinct roles in ***D. melanogaster***, where the former is involved in microRNA (miRNA) biogenesis while the latter participates in dsRNAs processing into small-interfering RNAs (siRNAs) (Lee et al., 2004). Phylogenetic and sequence analysis of malacostracan Dicer proteins revealed that they form two clusters representing Dicer-1 and Dicer-2 (fig. 9B). With a few exceptions, most malacostracans have single-copy homologs of Dicer-1 and Dicer-2 proteins; the krill species ***E. superba*** and ***M. norvegica*** have only Dicer-2 transcripts (supplementary table 8). With respect to Argonautes, we observed that most malacostracan species have multiple copies of this gene. We show that these putative Argonautes form a separate cluster from the closely related Piwi proteins (supplementary figure 7). We identified single-copy homologs of Argonaute-1 and multiple copies of Argonaute-2 in malacostracans (supplementary figure 7). Duplications of Argonaute-2 have occurred independently in specific lineages because variable copy number of this protein is reported in chelicerates but not in insects (Schnettler et al., 2014; Palmer et al., 2015). Also, the longer branch lengths of Argonaute-2 proteins indicate that sequence evolution is higher than those of Argonaute-1 (supplementary figure 7). In ***L. vannamei***, only Argonaute-2 is responsive to dsRNA (Labreuche et al., 2010). Hence, it was thought that Argonaute-1 operates through the miRNA pathway in shrimps (Chen et al., 2012). As in arthropods, the miRNA pathway is associated with crustacean antiviral defence (Lu et al., 2009; Fullaondo et al., 2012). The expression of miRNAs in ***M. japonicus*** was differentially regulated upon viral challenge (Ruan et al., 2011) and in other systems, viral infection results in the modification of host miRNA profiles (Li et al., 2011; Du et al., 2011; Tian et al., 2012). Components of miRNA biogenesis are intact in malacostracans; we identified single-copy homologs of ***Drosha*** and ***Pasha*** (fig. 9C and E). Research in the area of RNAi-mediated antiviral immunity has remained comparatively sparse in crustaceans, despite its rich therapeutic potential. Our results provide independent evidence that malacostracans have a naturally occurring antiviral defence mechanism in place. Much more needs to be done to understand the role of RNAi in innate immunity before it can be exploited for host defence against viral infections.

**Figure 9.**
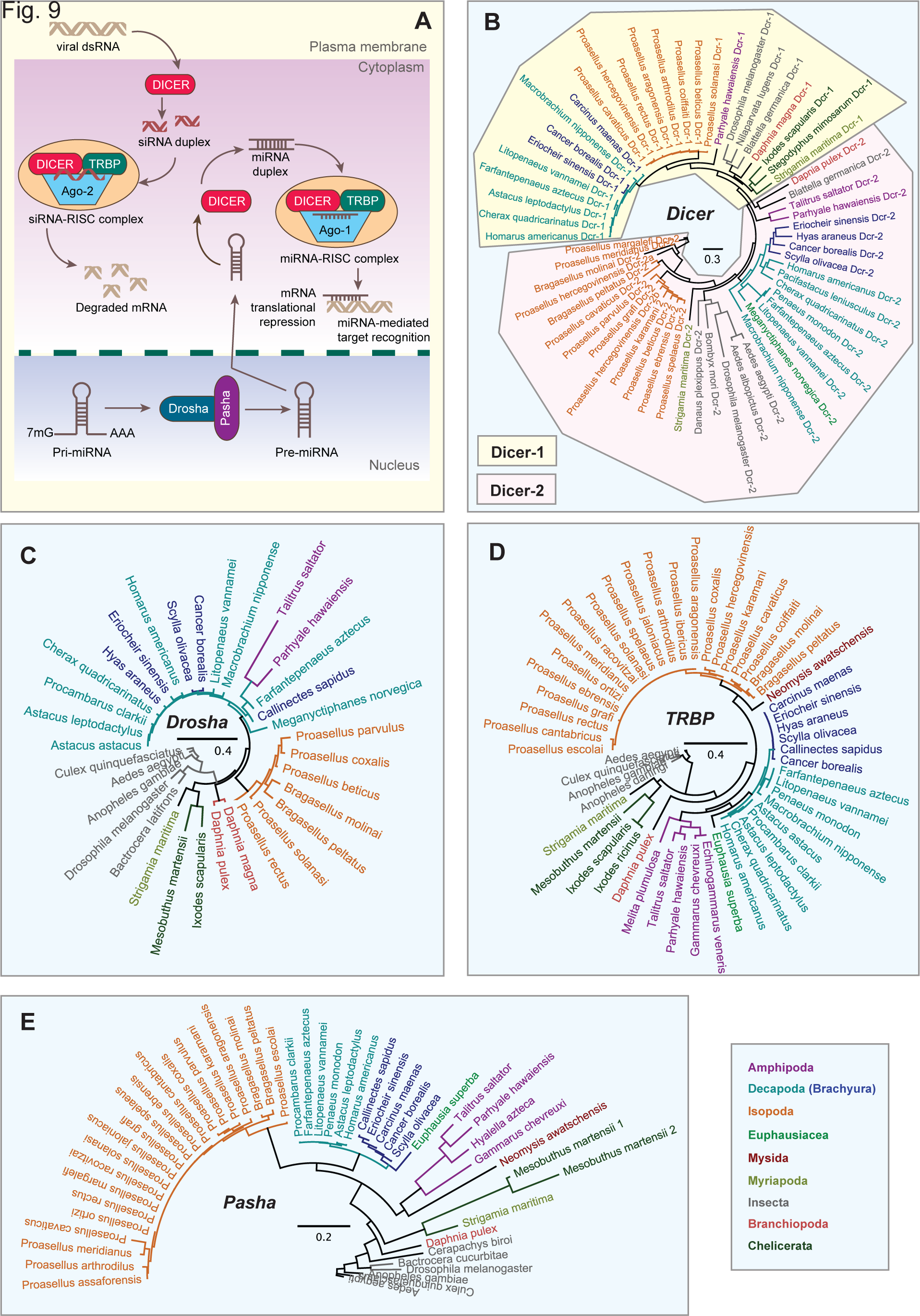
RNA interference (RNAi) pathway members in malacostracans. **(A)** Exogenous viral dsRNA is imported into the cytoplasm and is cleaved by the endoribonuclease from the RNase III family, Dicer. Cleaved fragments of dsRNA are known as small interfering RNAs (siRNAs). Dicer contains both Piwi/Argonaute/Zwille (PAZ) and helicase domains. Dicer activates the RNA-induced silencing complex (RISC), which is comprised of Argonaute-2 and the transactivating response RNA-binding protein (TRBP). Single-stranded siRNAs are incorporated into a RISC complex, upon which the siRNAs form complementary base-pairing to target mRNA and mRNA cleavage ensues. Other RISC-associated proteins include R2D2 and Loquacious in ***D. melanogaster.*** Dicer is also involved in microRNA (miRNA) biogenesis. Encoded by the genome, miRNAs are involved in the regulation of gene expression in the RNAi pathway. Transcribed by RNA polymerase II, Pri-miRNA is a long primary transcript of miRNAs and is processed into a stem-loop containing pre-miRNA by the microprocessor complex consisting of a ribonuclease III enzyme Drosha and partner of Drosha (Pasha), which is also known as DGCR8, protein. Pre-miRNA enters the cytoplasm and is cleaved by Dicer to generate a mature miRNA that is then integrated into the RISC complex. The miRNA-targeted transcript is either degraded or silenced. Phylogenetic trees of **(B)** Dicer, **(C)** Drosha, **(D)** TRBP and **(E)** Pasha are constructed using the maximum-likelihood method from an amino acid multiple sequence alignment. Taxa labels are depicted as their respective colour codes. Bootstrap support values (n=1000) for all trees can be found in supplementary figure 8. Scale bar represents substitution per site.

### Four novel gene families with potential involvement in malacostracan immunity

We have shown that although most canonical immunity genes and pathways in malacostracans share broad conservation with arthropods, lineage specific diversifications and gene duplications are common, which together suggests that lineage specific immune components may exist. With the advent of high-throughput sequencing, we are now able to tap into the availability of growing transcriptomic resources to find currently unknown proteins that might have potential involvement in host defence. Here, we present four novel gene families classified on the basis of shared domains implicated in immune function. We obtained a set of crustacean specific proteins from an orthology analysis using complete arthropod genomes (Kao et al., 2016). This list contained 750 protein sequences that have no significant blast hit to any other sequences in the NCBI nr database. We filtered this list down to 82 genes based on the presence of known Pfam domains (Bateman et al. 2004) and then down to a selection of 4 genes with domains suggestive of immune function (supplementary table 9). We used ***P. hawaiensis*** as a starting point for this analysis as this is the only complete Malacostraca genome available to date. While we are aware of potential limitations of this approach; for example, we may miss gene families that are not present in ***P. hawaiensis***, since we were interested in genes that are found across all five malacostracan orders, we were able to rationalise the use of ***P. hawaiensis***, an established to laboratory organism, as a basis for comparison. Considering genes with known Pfam domains allowed us to investigate potential structural characteristics that have implied immune function. We observed broad conservation of all four novel gene families in Malacostraca. We have named these gene families, pending functional studies, according to their Pfam annotations: 1) chitin binding peritrophin-A family, 2) death domain family, 3) ML domain family and 4) von Willebrand factor type A family (supplementary table 9). We found that these genes are all expressed in the ***P. hawaiensis*** and that they exhibited differential expression patterns between developmental stages and tissue types (fig. 10; supplementary table 10).

**Figure 10.**
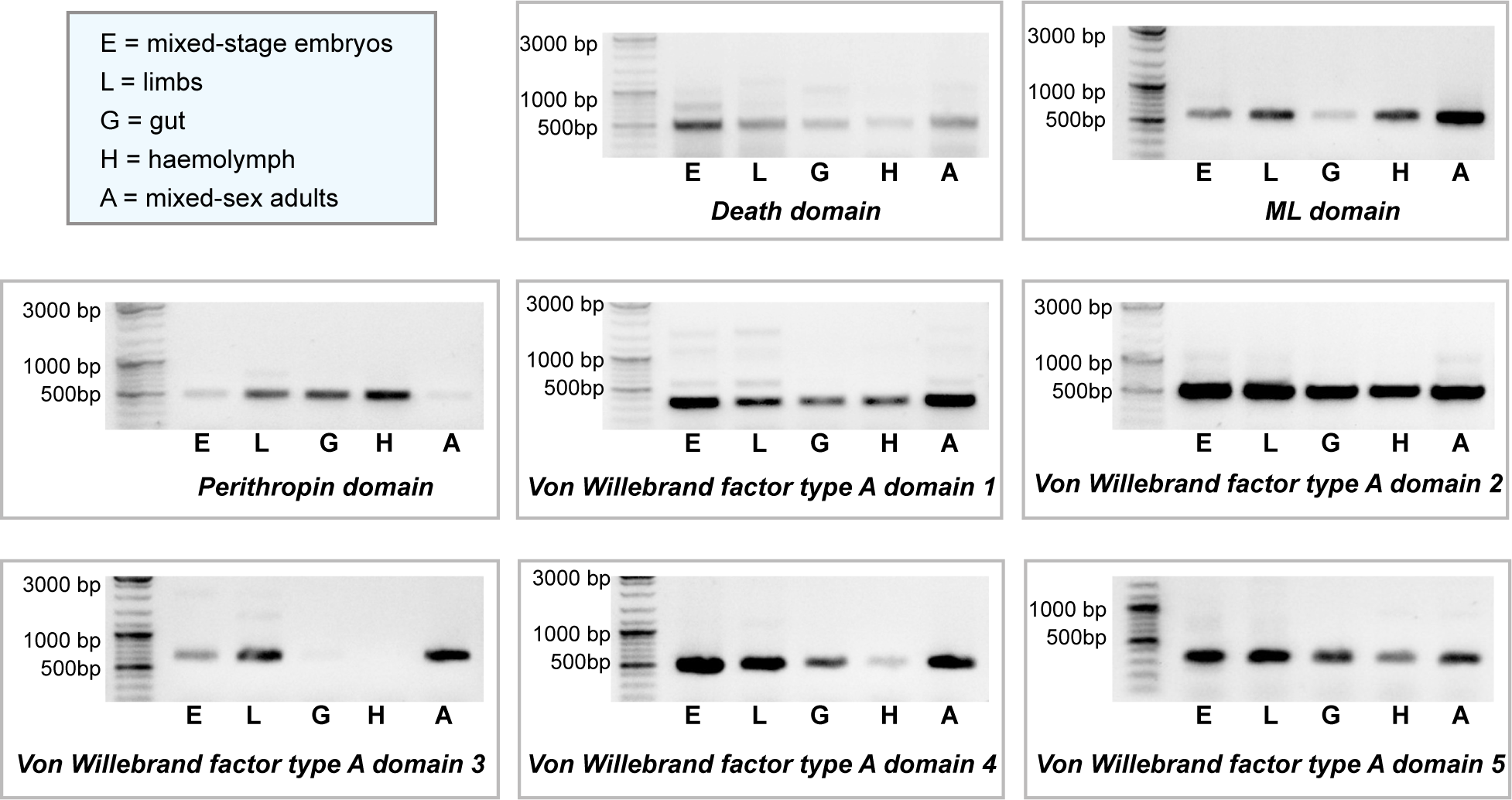
RT-PCR results showing expression patterns for eight novel malacostracan genes in five ***Parhyale hawaiensis*** tissue samples. These genes represent the four novel malacostracan gene families with potential function in innate immunity; genes containing the death domain, ML domain, peritrophin domain and the von Willebrand factor type A domain. Tissue samples were obtained from ***P. hawaiensis:*** mixed stage embryos, limbs, gut, hemolymph and mixed-sex mature animals.

Peritrophins are chitin-binding proteins originally isolated from the insect gut peritrophic membrane (Loongyai et al., 2007). Peritrophic membrane is thought to constitute a barrier for midgut epithelial cells to prevent the entry of microbes (Lehane, 1997; Wang and Granados, 1997; Terra, 2001). Peritrophin-like proteins are characterised by peritrophin domains. One example is the peritrophin-A domain, which contains six conserved cysteine residues separated by other nonconserved amino acids (fig. 11; Tellam et al., 1999). It has been shown recently that crustaceans also have peritrophin-like genes. Penaeid shrimps express peritrophins during oogenesis and these proteins have roles in the protection of spawned eggs against ***Vibrio*** (Khayat et al., 2001; Loongyai et al., 2007). A peritrophin-like gene cloned from ***Fenneropenaeus chinensis*** could bind chitin and Gram-negative bacteria (Du et al., 2006) and another peritrophin-like protein from ***Exopalaemon carinicauda*** is involved in WSSV infection (Wang et al., 2013). To date, only 10 peritrophin-like genes have been identified in crustaceans and no reports exist beyond decapod species (Khayat et al., 2001; Du et al., 2006; Chen et al., 2007; Loongyai et al., 2007; Chen et al., 2009; Wang et al., 2013; Huang et al., 2015). Here, we identified 80 novel peritrophin-like genes across five Malacostraca order (supplementary table 9; fig. 11A). This novel family of peritrophin-like proteins have no significant similarities to other peritrophin genes previously reported in crustaceans or arthropods. They share 18 conserved cysteine residues, a chitin-binding peritrophin-A domain and multiple conserved aromatic amino acids (fig. 11B). In ***P. hawaiensis***, we observed that one peritrophin-like gene is highly expressed in gut, hemocyte and limb samples but not in whole embryonic or adult tissue samples (fig. 10). This corroborates the observation that other peritrophins are found in the gut peritrophic membrane. They may serve additional roles in innate immunity since we also saw increased expression in circulating hemolymph (fig. 10).

**Figure 11.**
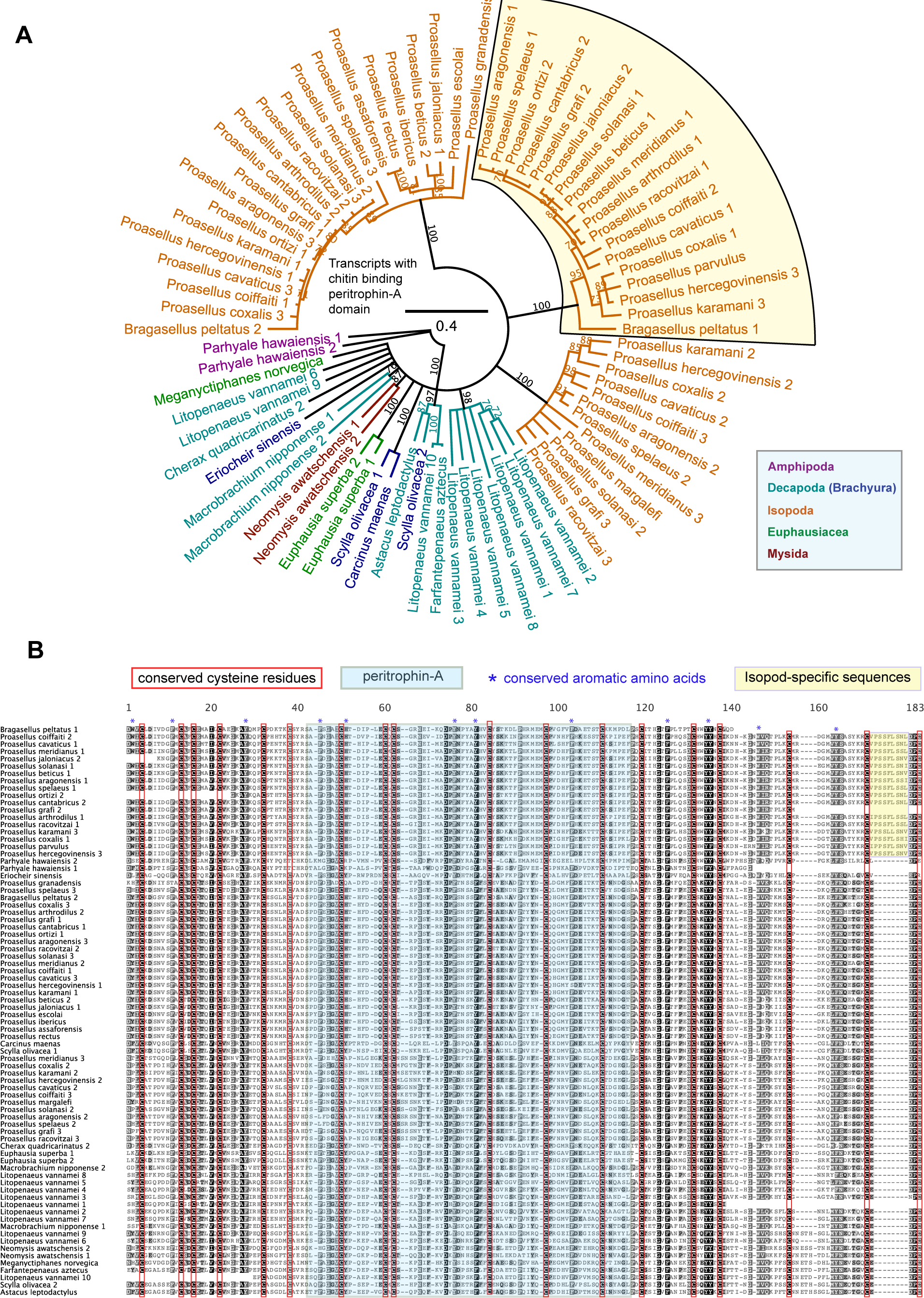
Novel gene family with potential innate immunity function characterised by the chitin binding peritrophin-A domain (pfam01607) found only in malacostracans. **(A)** Phylogenetic tree of proteins containing the chitin binding peritrophin-A domain is constructed using the maximum-likelihood method from an amino acid multiple sequence alignment. Taxa labels are depicted as their respective colour codes. Node labels represent bootstrap support values from 1000 replicates. Scale bar represents substitution per site. **(B)** Multiple sequence alignment of peritrophin-A proteins showing the conserved peritrophin-A domain marked with a blue box, conserved cysteine residues marked in red boxes and conserved aromatic amino acids indicated by blue asterisks. The yellow shaded box represents a group of proteins with additional isopod-specific sequences.

The second novel gene family specific to malacostracans is characterised by a death domain (DD). The DD superfamily represents evolutionarily conserved proteins of four subfamilies: the DD subfamily, the caspase recruitment domain (CARD) subfamily, the pyrin domain (PYD) subfamily and the death effector domain (DED) subfamily (Reed et al., 2004; McEntyre et al., 2004). Many DD superfamily members are involved in the regulation of immune response. Some examples are Imd, FADD and DREDD of the Imd pathway (Kischkel et al., 1995; Chinnaiyan et al., 1995; Muzio et al., 1996; Boldin et al., 1996; Georgel et al., 2001), MyD88, Tube and Pelle of the Toll pathway (Lemaitre et al., 1996; Sun et al., 2002) and receptor interacting protein (RIP), tumor necrosis factor receptor-1 (TNFR1), TNFR-associated death domain (TRADD) and (MAP kinase-activating death domain) MADD of the TNF pathway (Stanger et al., 1995; Hsu et al., 1995; Hsu et al., 1996a; Hsu et al., 1996b; Schievella et al., 1997). This new malacostracan DD gene family is a group of novel single-copy transcripts sharing a C-terminal DD (fig. 12A). This family is specific to Malacostraca and has no significant blast results to the nr database or any other DD-containing proteins (supplementary table 9). The N-terminal region of this family exhibited considerable similarities within members of this group but no known domains could be determined by HMM in this region (Fig. 12B). Experimental confirmation revealed that embryonic samples show the highest expression of a ***P. hawaiensis*** DD gene (fig. 10). This observation is intriguing because programmed cell death can be a key process during animal embryogenesis (Abrams et al., 1993) and many DD proteins are shown to be involved in apoptosis (Chinnaiyan et al., 1995; Nagata 1997).

**Figure 12.**
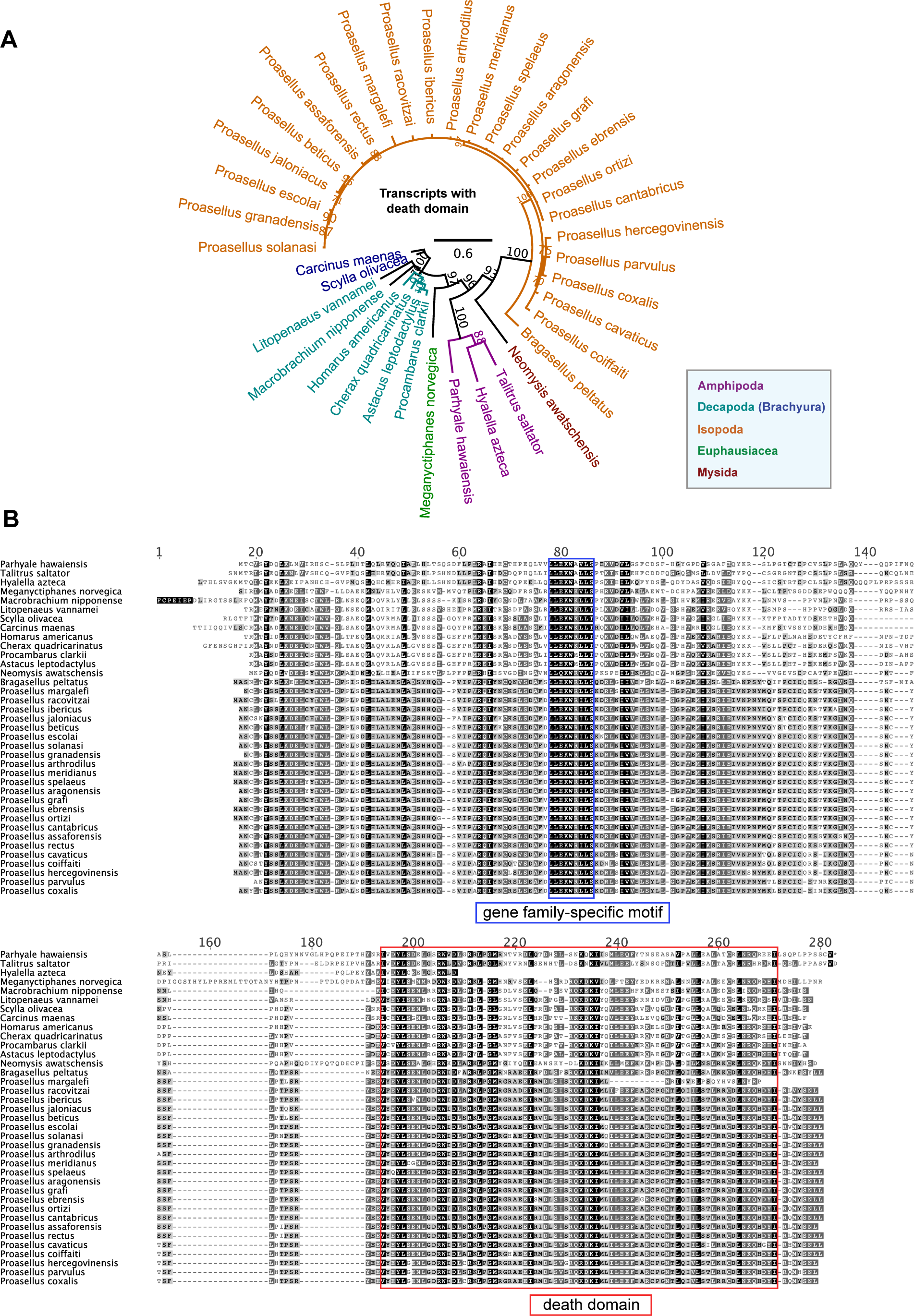
Novel gene family with potential innate immunity function characterised by the death domain (pfam00531) found only in malacostracans. **(A)** Phylogenetic tree of proteins containing the death domain is constructed using the maximum-likelihood method from an amino acid multiple sequence alignment. Taxa labels are depicted as their respective colour codes. Node labels represent bootstrap support values from 1000 replicates. Scale bar represents substitution per site. **(B)** Multiple sequence alignment of the death domain family showing the death domain marked with a red box and a novel gene-family-specific motif marked with a blue box.

The third novel gene family we identified in malacostracans is characterised by the presence of a MD-2-related lipid recognition (ML) domain. The ML domain was originally identified from a group of unknown proteins that share regions of homology with the MD-2 protein (Inohara et al., 2002). We identified 39 transcripts containing the ML domain; they have no significant blast hits to any known genes so we named this the malacostracan ML family (supplementary table 9). The malacostracan ML family contains six conserved cysteine residues and these residues may be involved in the formation of disulfide bonds (fig. 13B). Mutation of a conserved cysteine in MD-2 abolishes the response to LPS, which suggests a role of the disulfide bond in MD-2 function (Schromm et al., 2001). Members of this ML family are expressed in a wide range of tissue types; we observed expression in brain and nervous system samples (fig. 13A; supplementary table 9). ML genes in ***P. hawaiensis*** exhibited the highest level of expression in adult tissue samples and to lesser degrees in embryos or gut samples (fig. 10). Since ML proteins have been implicated in lipid signalling, metabolism and immunity (Kirchhoff et al., 1996; Okamura et al., 1999; Naureckiene et al., 2000; Inohara et al. 2002), increased expression of ***P. hawaiensis*** ML in whole adult samples may imply a metabolic role in addition to host defence, as expression is also be observed in hemocytes. Although not much is known about the direct roles of ML proteins in immunity, others have proposed that they could participate as lipid-binding cofactors in the recognition of pathogenic agents (Inohara et al., 2002). In mammals, MD-2 directly binds LPS through the MD-2-TLR4 complex of the Toll signalling pathway (Shimazu et al., 1999; Viriyakosol et al., 2001; Gangloff et al., 2004). MD-2 has a leader sequence for endoplasmic reticulum targeting and secretion but lacks any transmembrane domain (Visintin et al., 2001). We predicted the presence of putative N-terminal transmembrane topologies in the malacostracans ML family, which indicates that they could be anchored to the cell membrane and have unknown PRR functions rather than being secreted (fig. 13B).

**Figure 13.**
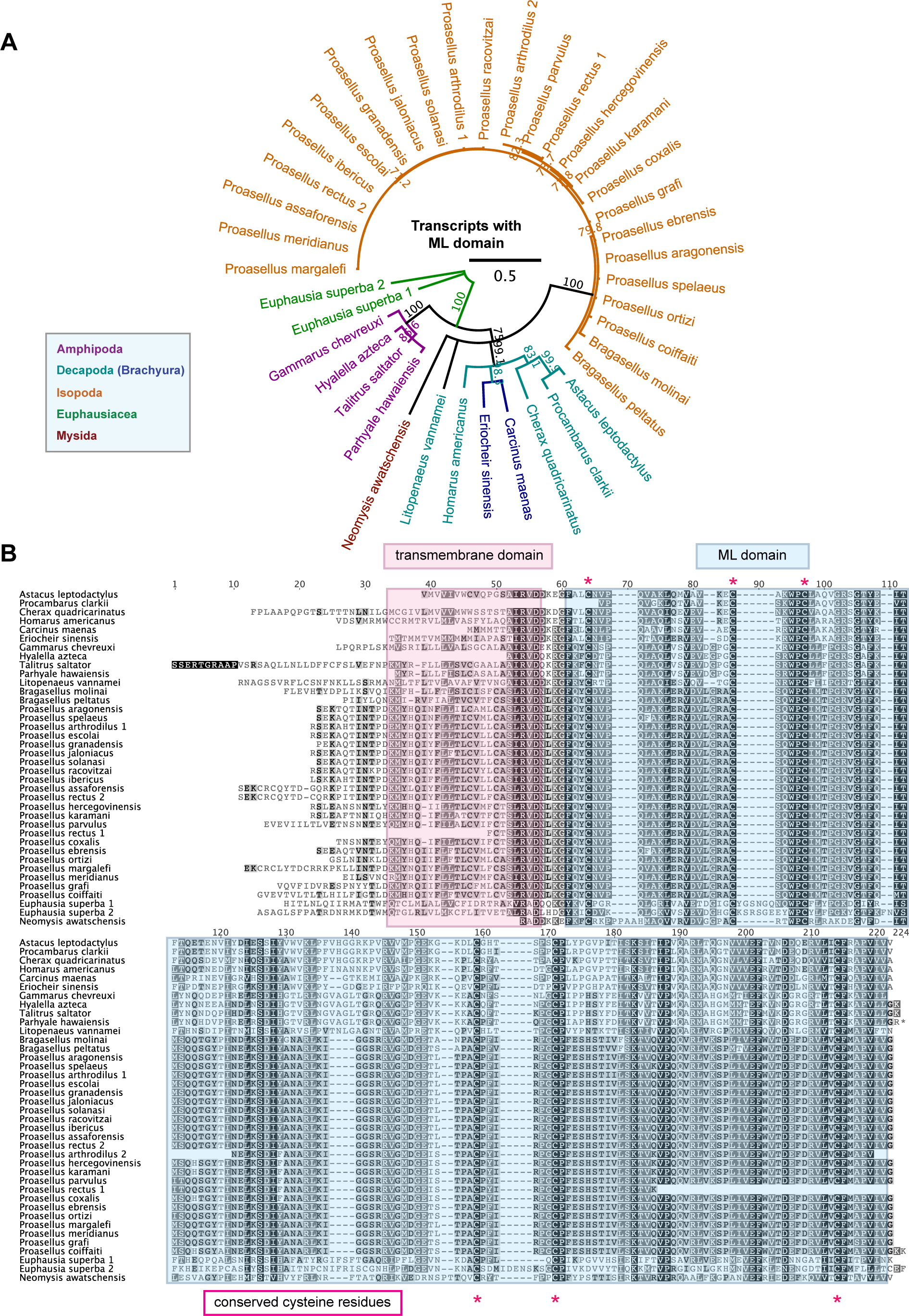
Novel gene family with potential innate immunity function characterised by the ML domain (pfam02221) found only in malacostracans. **(A)** Phylogenetic tree of proteins containing the ML domain is constructed using the maximum-likelihood method from an amino acid multiple sequence alignment. Taxa labels are depicted as their respective colour codes. Node labels represent bootstrap support values from 1000 replicates. Scale bar represents substitution per site. **(B)** Multiple sequence alignment of the ML family showing the ML domain characterised by six conserved cysteine residues marked with red asterisks. Transmembrane domains were predicted using the Phobius program (Käll et al., 2004; Käll et al., 2007) and are annotated with a pink box.

The final family of novel malacostracan genes with potential involvement in immunity has the von Willebrand factor type A (VWA) domain. Von Willebrand factor (VWF) proteins were discovered in patients with blood clotting disorders, named the von Willebrand disease (Alexander et al., 1953; Quick et al., 1953). They are ubiquitous in blood plasma and connective tissues and have functions in binding blood clotting factors and in mediating platelet adhesion at the site of vascular injury (Houdijk et al., 1985; Marchese et al., 1995; Wu et al., 1996). It was thought that enzymatic components of the blood clotting and complement system utilise similar macromolecular building blocks that existed before the protostome-deuterostome divergence (Krem et al., 2002). Since arthropods have open circulatory systems and lack adaptive immunity, hemolymph clotting is a pivotal part of the immune response because it functions not only to prevent hemolymph loss but also to immobilise pathogens at the site of wound (Opal, 2000). Arthropod hemolymph is also sensitive to small amounts of microbial polysaccharides (Iwanaga et al., 1998, Armstrong et al., 2003; Isakova et al., 2003) and clotting enzymes in arthropods emerged via convergent evolution from the assembly of domains found in vertebrate factors rather than being exact orthologs (Krem et al., 2002). Here, we describe the novel observation of a gene family that contained a VWA domain (supplementary table 9). They are present as multiple-copy homologs; we identified 87 genes from all five malacostracan orders and they have no significant sequence homology to any other known sequences (fig. 14). The malacostracans VWA family is made up of members with long transcripts up to 7kb in length, with coding sequences translated to polypeptides of up to 1500 amino acids; this feature is commonly seen in other VWF proteins (Verweij et al., 1986; Sadler, 1998). We observed expression in tissue specific datasets obtained from the brain, nervous system, hepatopancreas, hemocytes, gill and eyestalk (supplementary table 9). From our RT-PCR results, the five VWA genes in ***P. hawaiensis*** exhibited diverse patterns of expression; e.g. the VWA2 gene appears to be highly expressed in embryos, limbs, gut, hemolymph and adult tissues while the VWA3 gene is not expressed in gut and hemolymph samples (fig. 10). In summary, we have described four novel gene families specific only to malacostracans, which are good candidates for fulfilling roles in host defence. These genes offer new avenues for research and further analysis will be required to ascertain if they fit into the conceptual framework of innate immunity. Given our conservative approach to identifying gene families with domains that relate to known Pfam domains, it seems likely further comparative study, coupled with functional genomics and immunobiology approaches, will identify more malacostracan specific immune related genes.

**Figure 14.**
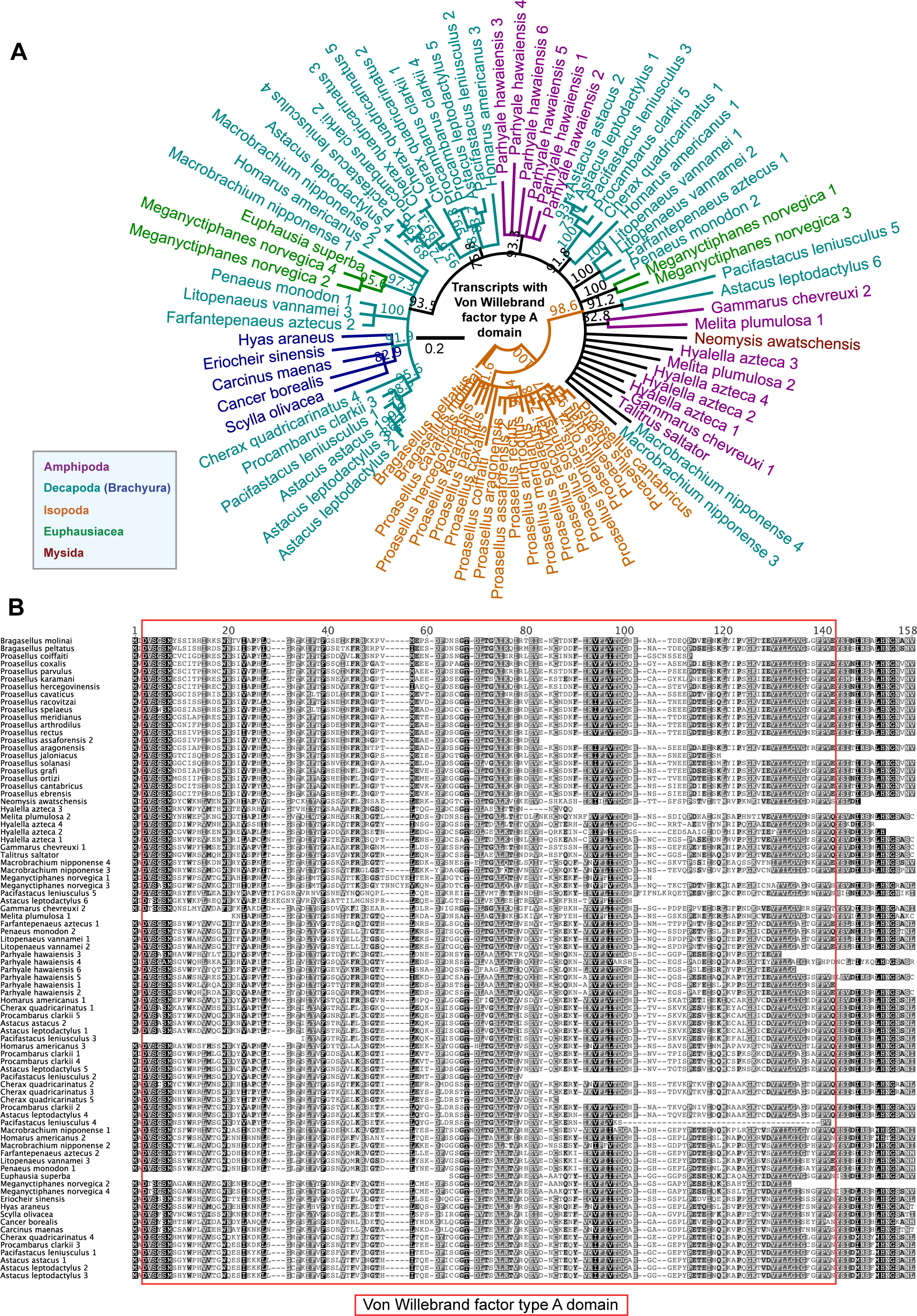
Novel gene family with potential innate immunity function characterised by the Von Willebrand factor type A domain (pfam05762) found only in malacostracans. **(A)** Phylogenetic tree of proteins containing the Von Willebrand factor type A domain is constructed using the maximum-likelihood method from an amino acid multiple sequence alignment. Taxa labels are depicted as their respective colour codes. Node labels represent bootstrap support values from 1000 replicates. Scale bar represents substitution per site. **(B)** Multiple sequence alignment of the Von Willebrand factor type A domain from this gene family.

## Conclusion

The recent availability of transcriptome sequences of distantly related malacostracan species has allowed us to describe molecular components of their innate immune systems at a new level of detail. This data is now available to the community to inform the next stages of immune research to underpin important aquaculture developments. By separating the immune response into successive phases, we observed dynamic evolutionary adaptations in the pathogen recognition phase, signal transduction and effector response systems.

Malacostracans achieve flexibility in recognising infections through the divergent evolution of certain PRR families, notably the gene expansions of GNBPs and CTLs. Upon recognition, several enzymatic cascades are involved in signal modulation and these too have novel evolutionary features. Malacostracans achieve diversity in modulation components through gene duplications of modulatory families involving CLIP serine proteases and Spätzle. When drawing comparisons to other arthropods, we observed novelties in these immune modulation components and are able to form strong evolutionary hypotheses as when key pathways evolved or diverged (e.g. the invention of proPO at the base of Pancrustacea). Core immune signal transduction pathways are largely conserved in malacostracans, although several components of the Imd pathways have been lost. The Imd pathway is activated through the digestion of PGNs by PGRPs in ***D. melanogaster*** (Ferrandon et al. 2007).

PGRPs are previously thought to be lost in Crustacea; as ***D. pulex*** lack PGRPs (McTaggart et al. 2009) and no PGRPs are present in the ***P. hawaiensis*** genome (Kao et al. 2016). Despite this being true for most malacostracan datasets we considered here, we were able to identify four PGRP genes spread across the Amphipoda, Isopoda and Decapoda orders, which are predicted to be catalytically active based on the presence of essential residues (supplementary figure 5A and D). Although unlikely, PGRPs in these species may have appeared through convergent evolution. Future studies will be required to determine the whether the biological roles of these PGRPs have any relevance in host defence and the evolutionary events that explain their relatively patchy phylogenetic distribution.

Effector mechanisms in malacostracans, like in other arthropods, are highly divergent and lineage specific. We described two malacostracan-specific AMPs previously confined to the Decapoda, and show that members of these families are widespread in other non-decapod malacostracan species. Crustaceans are regularly exposed to viral components in their natural environments (Loh et al., 1997; Fuhrman, 1999; Liu et al., 2009) and hence need antiviral mechanisms in place to counteract infection. We demonstrate that malacostracans have intact siRNA and miRNA components.

Finally, we present four novel gene families in Malacostraca as potential key players of the innate immunity. We only addressed the structural significance of these genes in the context of host defence based on comparisons with other immune proteins containing similar structural features. More functional studies will be required in the future to ascertain the roles of these genes and their potential function in innate immunity before they can be confirmed as crustacean immune system components.

Resources presented in this study facilitate and expand the scope of both basic and applied research, in particular analyses on the mechanistic links between specific immune modules and overall host defence. Importantly our data suggest that non-decapod species, like the laboratory model ***P. hawaiensis***, may nonetheless be suitable for studying malacostracan specific immune mechanisms relevant to food crop species.

## Acknowledgments and funding information

This work was supported by the Biotechnology and Biological Sciences Research Council (grant number BBK0075641 to A.A.A.); Human Frontier Science Program Postdoctoral Fellowship (to A.G.L.) and the Elizabeth Hannah Jenkinson Fund (to A.G.L.). We would also like to thank Alessia Di Donfrancesco for her help on *P. hawaiensis* RNA extractions.

## Materials and Methods

### Innate immunity datasets and query sets

We retrieved complete transcriptome datasets for 55 malacostracan species from the European Nucleotide Archive (http://www.ebi.ac.uk/ena). These transcriptomes included those generated from specific tissue types or developmental stages. We also included the *Parhyale hawaiensis* transcriptome generated by a separate study (Kao et al., 2016). All analyses were performed on a total of 69 transcriptome datasets. A full list of datasets and accessions used in this study is listed in supplementary table 1. For query sequences used in homology searches, we retrieved a set of insect immunity genes from ImmunoDB (http://cegg.unige.ch/Insecta/immunodb, Waterhouse et al., 2007) and known malacostracan entries compiled from Uniprot and GenBank. Both gene sets are consolidated to generate a core set of protein sequences used as queries.

### Identification of Malacostraca innate immunity genes

To generate a redundant set of malacostracan immunity gene orthologs, we used multiple Basic Local Alignment Search Tool (BLAST)-based approaches such as PSI-BLAST, BLASTp and tBLASTn with varying Blocks Substitution matrices. This redundant list was filtered by e-value of less than 10^−6^ and best reciprocal BLAST hits against the GenBank non-redundant (nr) database. We then filtered the best reciprocal nr BLAST hits by alignment length and identity (95%) to account for the potential of alleles assembling independently in polymorphic transcriptomes, the presence of conserved domains reported to be essential for function, and tree-based approaches to compile a final non-redundant list of Malacostraca innate immunity orthologs.

### Identification of conserved domains and phylogenetic tree construction

Malacostracan transcripts were in silico translated according to the longest open-reading frames into protein sequences. Conserved domains of the malacostracan hits were annotated using the Batch CD-Search Tool by NCBI (https://www.ncbi.nlm.nih.gov/Structure/bwrpsb/bwrpsb.cgi). Hits without essential domains were discarded. Multiple sequence alignments of protein sequences were performed using MAFFT (Katoh et al., 2009). Phylogenetic trees were built from the MAFFT alignments using the WAG+G model in RAxML (Stamatakis, 2014) to generate best-scoring maximum likelihood trees. Multiple sequence alignment images were generated using the Geneious programme (Kearse et al., 2012). Graphical representations of Newick trees were also generated using Geneious.

### Identification of novel genes with putative immune function in Malacostraca

We previously performed orthologous group analyses in a separate study using complete arthropod genomes, which included the genome of the Malacostraca *P. hawaiensis* (Kao et al., 2016). We retrieved a list containing 750 protein sequences that were found only in *P. hawaiensis* and have no significant blast hits to any other sequences in the NCBI nr database (Kao et al., 2016). We performed a scan for the presence of Pfam domains (Bateman et al., 2004) using HMMER (http://hmmer.org/; Eddy, 2011; Finn et al., 2011) on these 750 sequences and identified 82 genes containing Pfam domains. Further examination of predicted Pfam domains revealed four genes with domains suggestive of immune function: 1) chitin binding peritrophin-A domain pfam01607, 2) death domain pfam00531, 3) von Willebrand factor type A domain pfam05762 and 4) ML domain pfam02221 (Supplementary table 9). These four genes were used as query sequences for Blast against the other malacostracan transcriptomes. We scanned putative malacostracan orthologues for transmembrane domains and signal peptides using the Phobius tool (Käll et al., 2004; Käll et al., 2007).

### RNA extraction and reverse transcriptase-polymerase chain reaction (RT-PCR) of novel Malacostraca immunity genes in *Parhyale hawaiensis*

Five types *P. hawaiensis* tissue samples were collected from: 1) 100 mixed-stage embryos, 2) amputated limb fragments from 15 adult animals, 3) dissected gut tissues from 20 adult animals, 4) hemolymph from 50 adult animals and 5) adult whole tissues from two males and two females. Embryos were dissected from gravid females and rinsed with molecular grade water. Prior to tissue collection from the adults, animals were washed with filtered artificial seawater followed by treating with a mixture of clove oil (Sigma) and milliQ water (1:5000 dilution) for anaesthetization. As soon as the animals stopped moving after several minutes, the clove oil mixture was rinsed off and the anaesthetized animals were rinsed with molecular grade water. Limb fragments were dissected using spring scissors (Fine Science Tools). Gut tissues were collected using a scalpel and fine forceps. Hemolymph samples were collected by allowing the animals to bleed out in molecular grade water. Immediately after collection, 1mL of Trizol reagent (Thermo Fisher Scientific) was added to the tissue samples in eppendorf tubes and samples were then snap frozen on dry ice. RNA extractions were performed according to the Trizol manufacturer's instructions. Concentrations of total RNA extracts were quantified using Qubit and Nanodrop. One microgram of total RNA from each tissue type was used for cDNA synthesis using the Qiagen QuantiTect Reverse Transcription Kit according to manufacturer's instructions. PCR on each gene was performed using Phusion High-fidelity polymerase (Thermo Fisher Scientific) and the following program was used for thermal cycling: 98C 30s, followed by 25 cycles of 98C 10s, 62C 30s, 72C 45s, and then 72C 5m. PCR products were ran on a 1% agarose gel and stained with SYBR Safe (Thermo Fisher Scientific). Primer sequences used for PCR are listed in supplementary table 10.

---

**supplementary figure 1.**
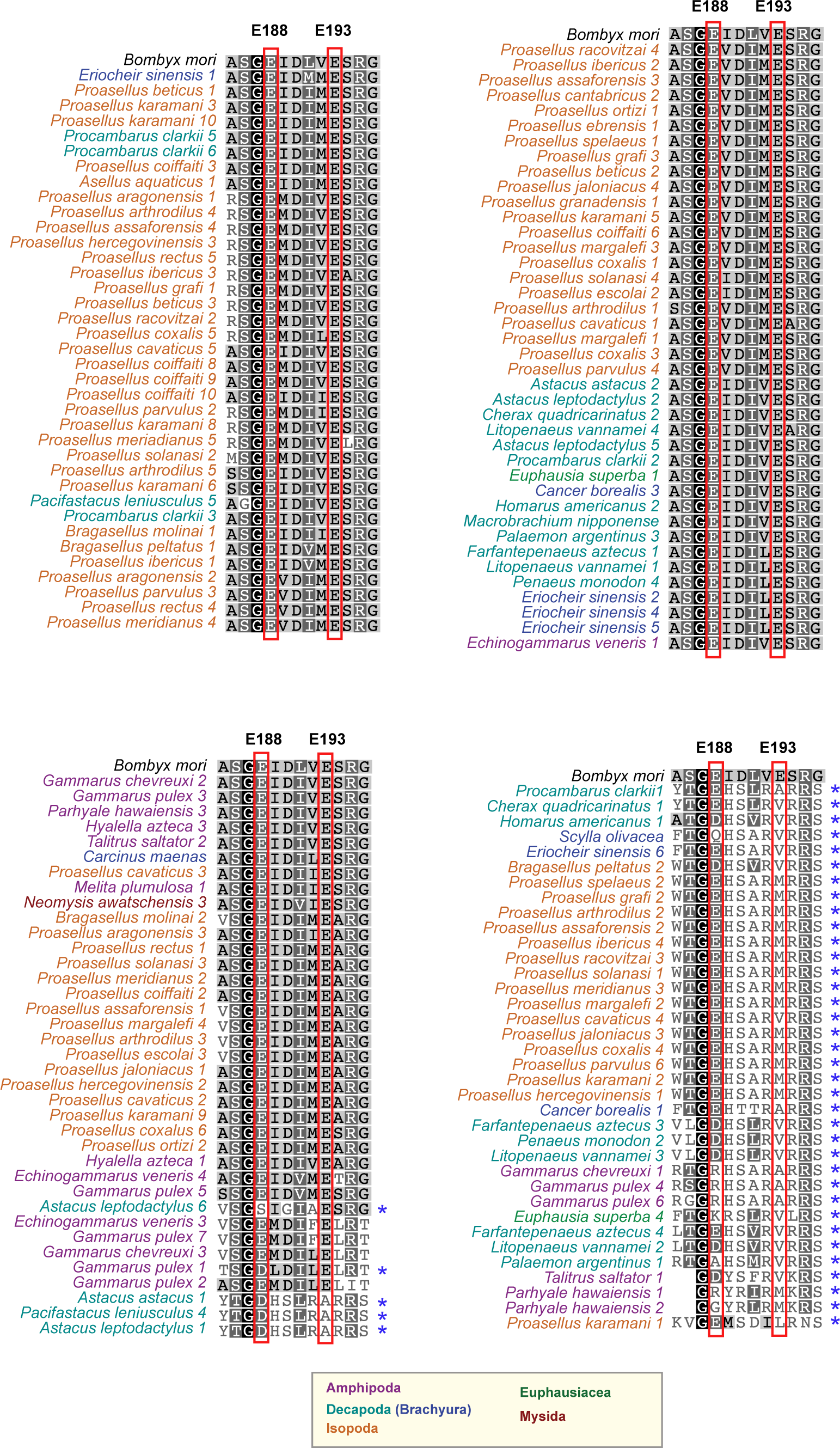
Multiple sequence alignment of the β-glucanase domains of Gram negative binding proteins of malacostracans together with the β-glucanase protein from *Bombyx mori* (NP_001159614.1). Two conserved Glu active site residues are labeled as E188 and E193 based on positions in the *B. mori* protein.

**supplementary figure 2.**
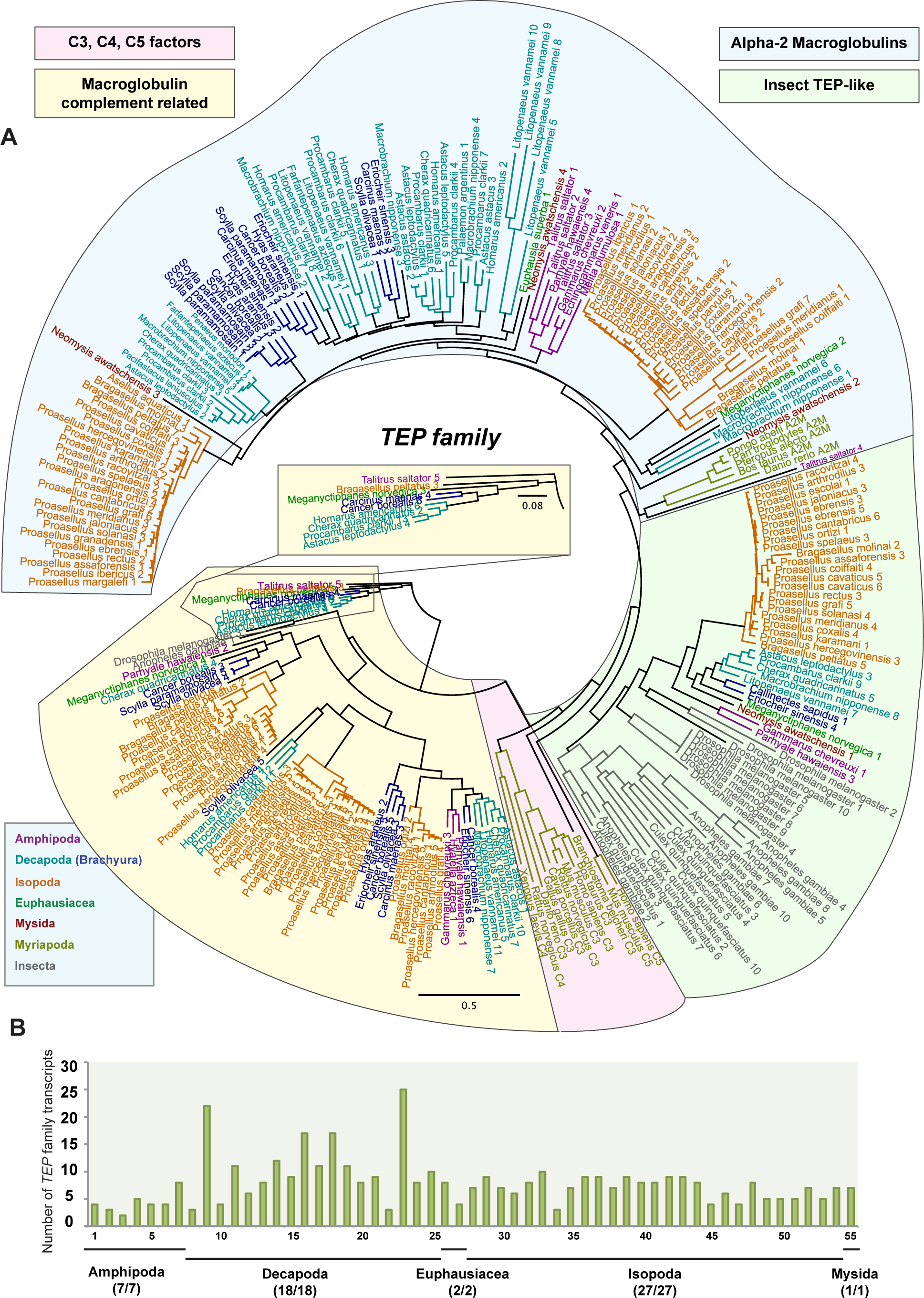
Thioester-containing protein (TEPs) family. **(A)** Phylogenetic tree of the TEP family is constructed using the maximum-likelihood method from an amino acid multiple sequence alignment. Amino acid sequences include the macroglobulin complement related proteins, the vertebrate C3, C4 and C5 complement factors, arthropod TEPs and the alpha-2 macroglobulin family. Taxa labels are depicted as their respective colour codes. Bootstrap support values (n=1000) for all trees can be found in supplementary figure 8. Scale bar represents substitution per site. **(B)** Graph of putative *TEP* family transcripts. The y-axes represent total number of genes identified in all 55 malacostracan species for each family. Each species is represented by a number on the X-axes and a complete list of species is available in Supplementary table 2. Black horizontal bars below each graph delimit the five orders of malacostracans and the numbers in parentheses (x/y) represent the following: x = number of species in which a particular gene family is found and y = total number of species in each order.

**supplementary figure 3.**
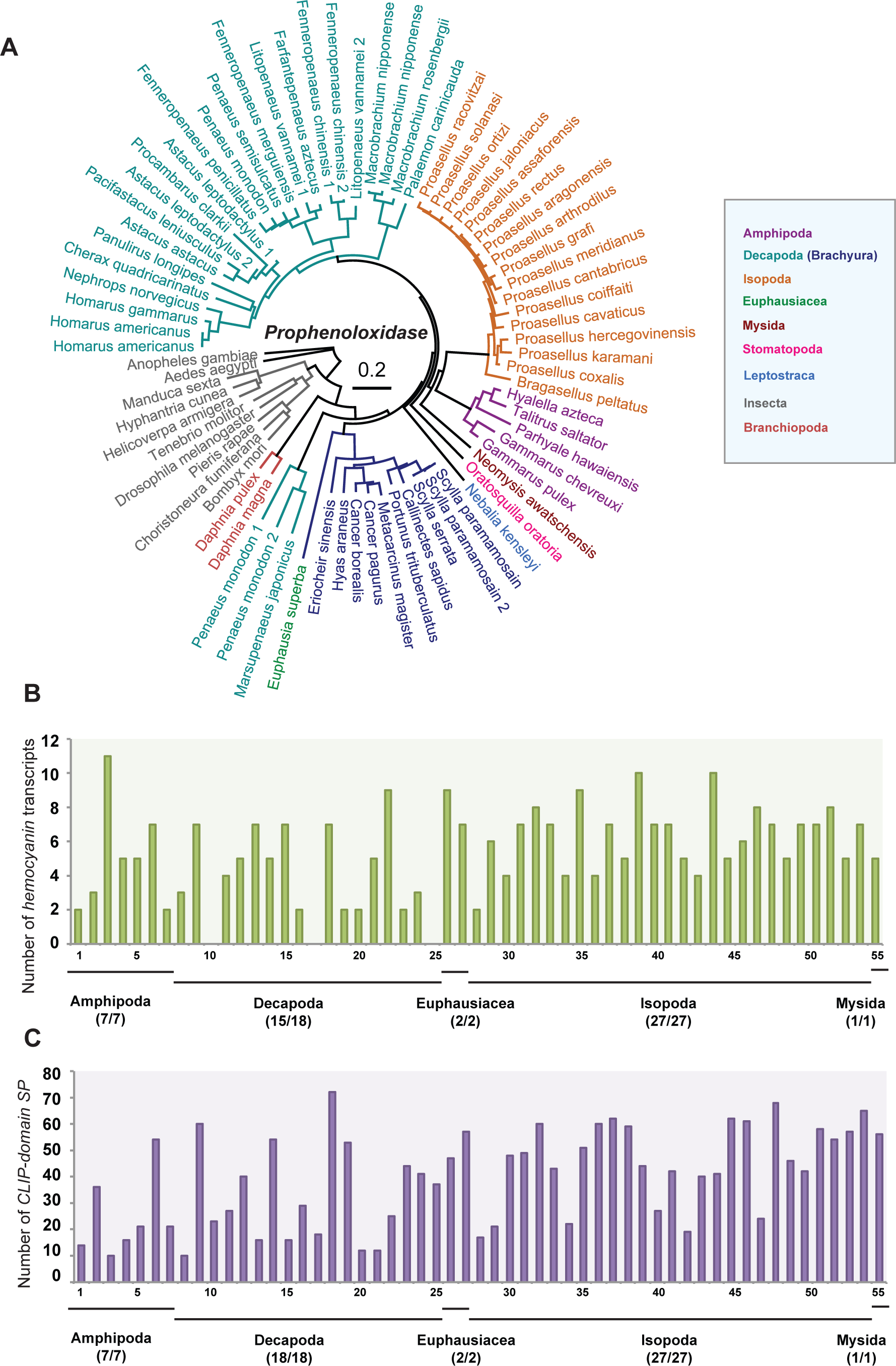
Prophenoloxidase activation system. **(A)** Phylogenetic tree of prophenoloxidase (proPO) is constructed using the maximum-likelihood method from an amino acid multiple sequence alignment. Taxa labels are depicted as their respective colour codes. Bootstrap support values (n=1000) for all trees can be found in supplementary figure 8. Scale bar represents substitution per site. The graphs represent the total number of **(B)** *hemocyanin* and **(C)** *CLIP-domain serine protease* transcripts in malacostracans. The y-axes represent total number of genes identified in all 55 malacostracan species for each family. Each species is represented by a number on the X-axes and a complete list of species is available in Supplementary table 2. Black horizontal bars below each graph delimit the five orders of malacostracans and the numbers in parentheses (x/y) represent the following: x = number of species in which a particular gene family is found and y = total number of species in each order.

**supplementary figure 4.**
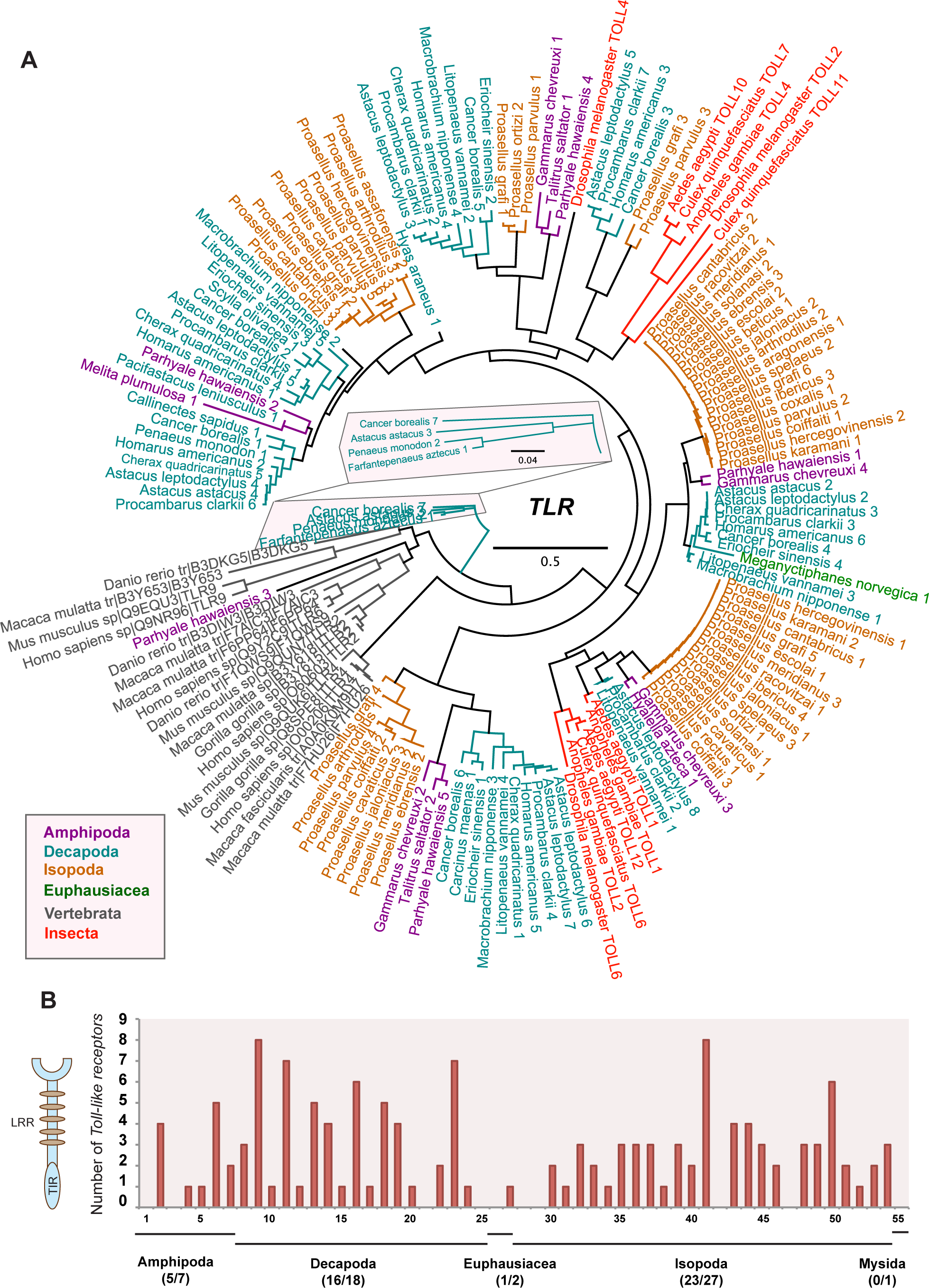
Malacostracans Toll-like receptors (TLRs). **(A)** Phylogenetic tree of TLRs is constructed using the maximum-likelihood method from an amino acid multiple sequence alignment of toll-IL-1 receptor (TIR) domains. Taxa labels are depicted as their respective colour codes. Bootstrap support values (n=1000) for all trees can be found in supplementary figure 8. Scale bar represents substitution per site. **(B)** Graph of putative *TLR* transcripts. The y-axes represent total number of genes identified in all 55 malacostracan species for each family. Each species is represented by a number on the X-axes and a complete list of species is available in Supplementary table 2. Black horizontal bars below each graph delimit the five orders of malacostracans and the numbers in parentheses (x/y) represent the following: x = number of species in which a particular gene family is found and y = total number of species in each order.

**supplementary figure 5.**
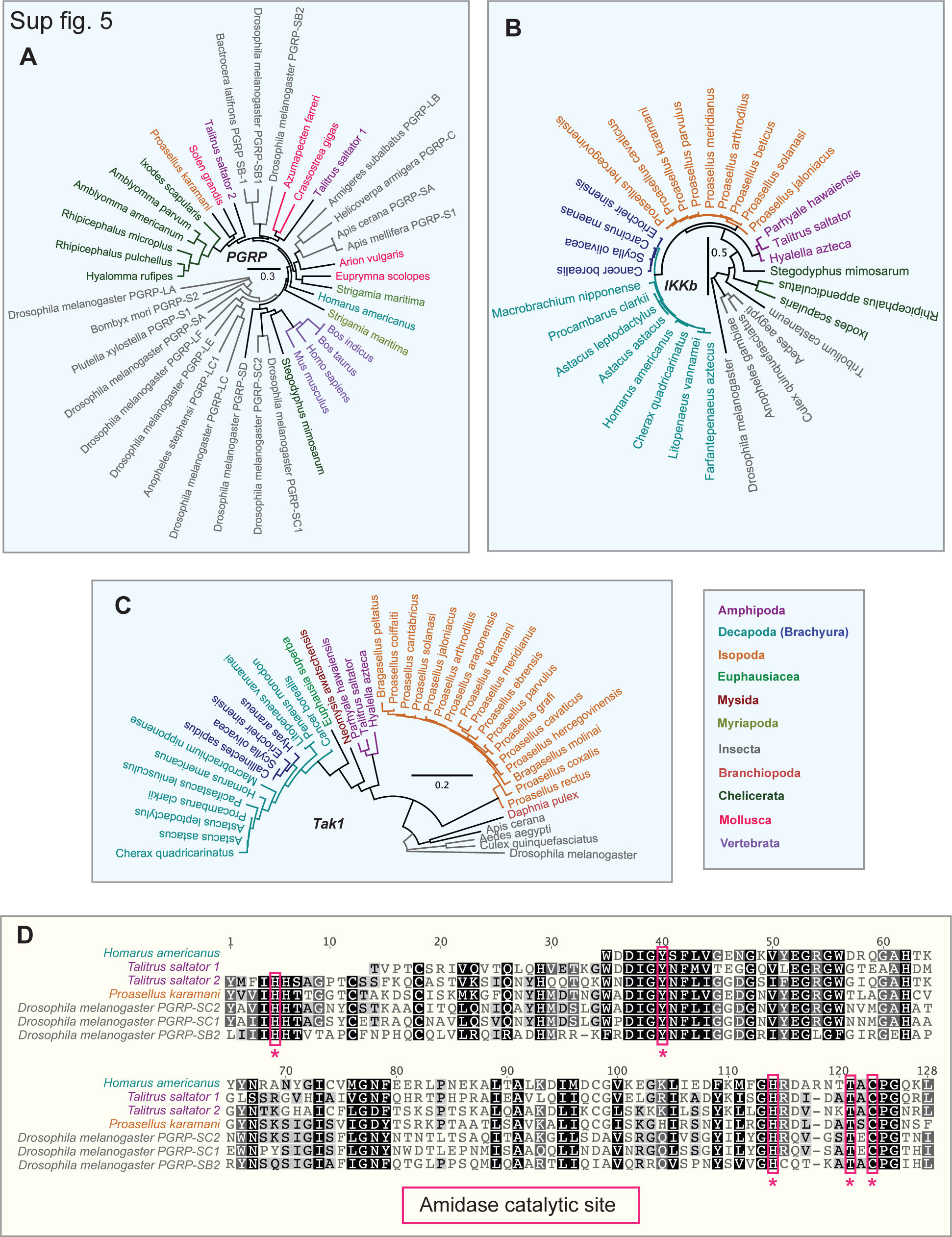
Additional members of the IMD pathway. Phylogenetic trees of **(A)** PGRP, **(B)** IκB kinase β (IKKb); also known as immune response deficient-5 (IRD-5) and **(C)** TAK1 are constructed using the maximum-likelihood method from an amino acid multiple sequence alignment. Taxa labels are depicted as their respective colour codes. Bootstrap support values (n=1000) for all trees can be found in supplementary figure 8. Scale bar represents substitution per site. **(D)** Multiple sequence alignment of four PGRPs identified in malacostracans. Conserved amidase catalytic residues are highlighted in red boxes.

**supplementary figure 6.**
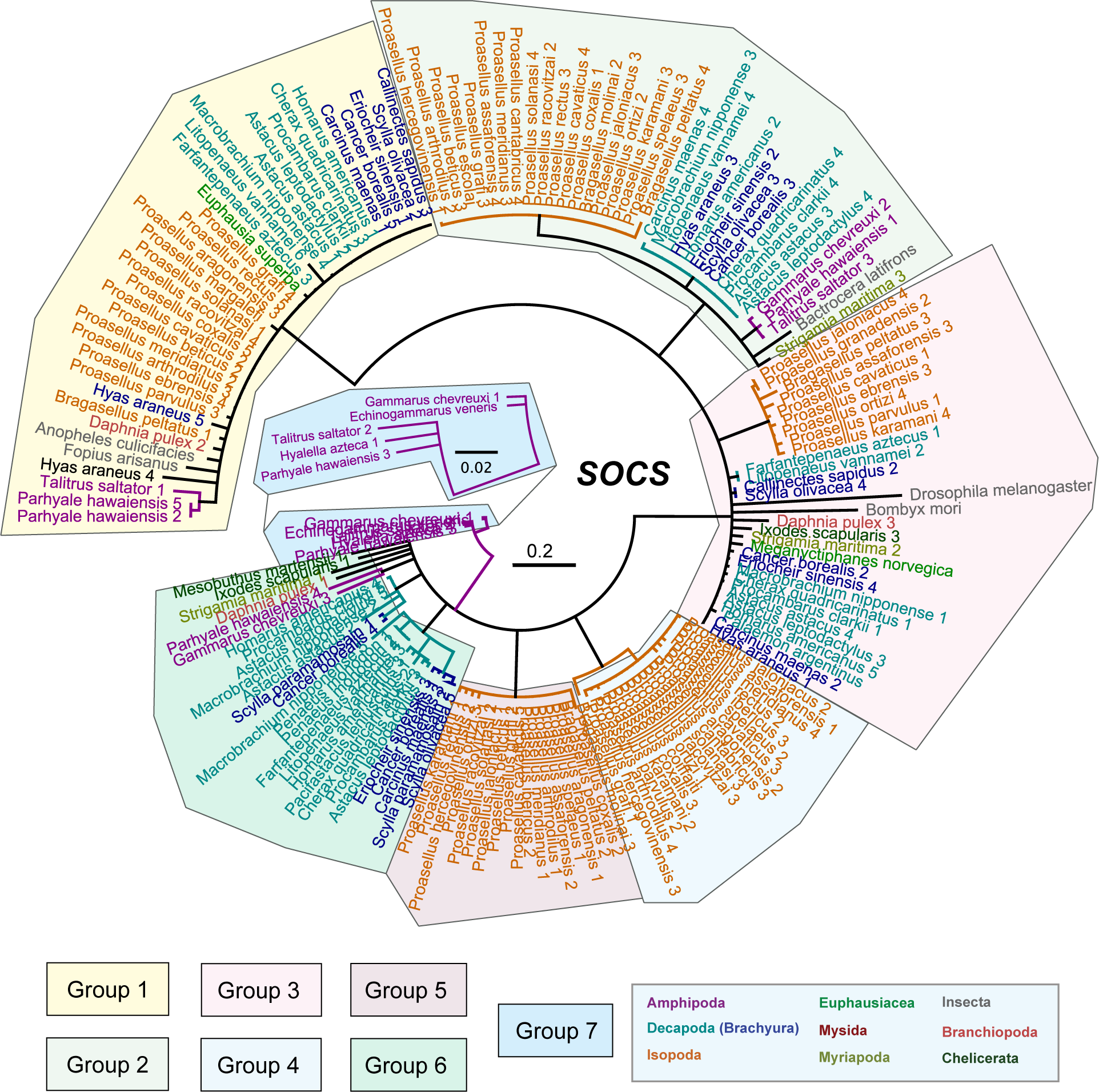
Suppressor of cytokine signalling (SOCS) gene family in malacostracans. Phylogenetic tree of the SOCS gene family is constructed using the maximum-likelihood method from an amino acid multiple sequence alignment. Seven main groups of SOCS proteins are identified in malacostracans. Taxa labels are depicted as their respective colour codes. Bootstrap support values (n=1000) for all trees can be found in supplementary figure 8. Scale bar represents substitution per site.

**supplementary figure 7.**
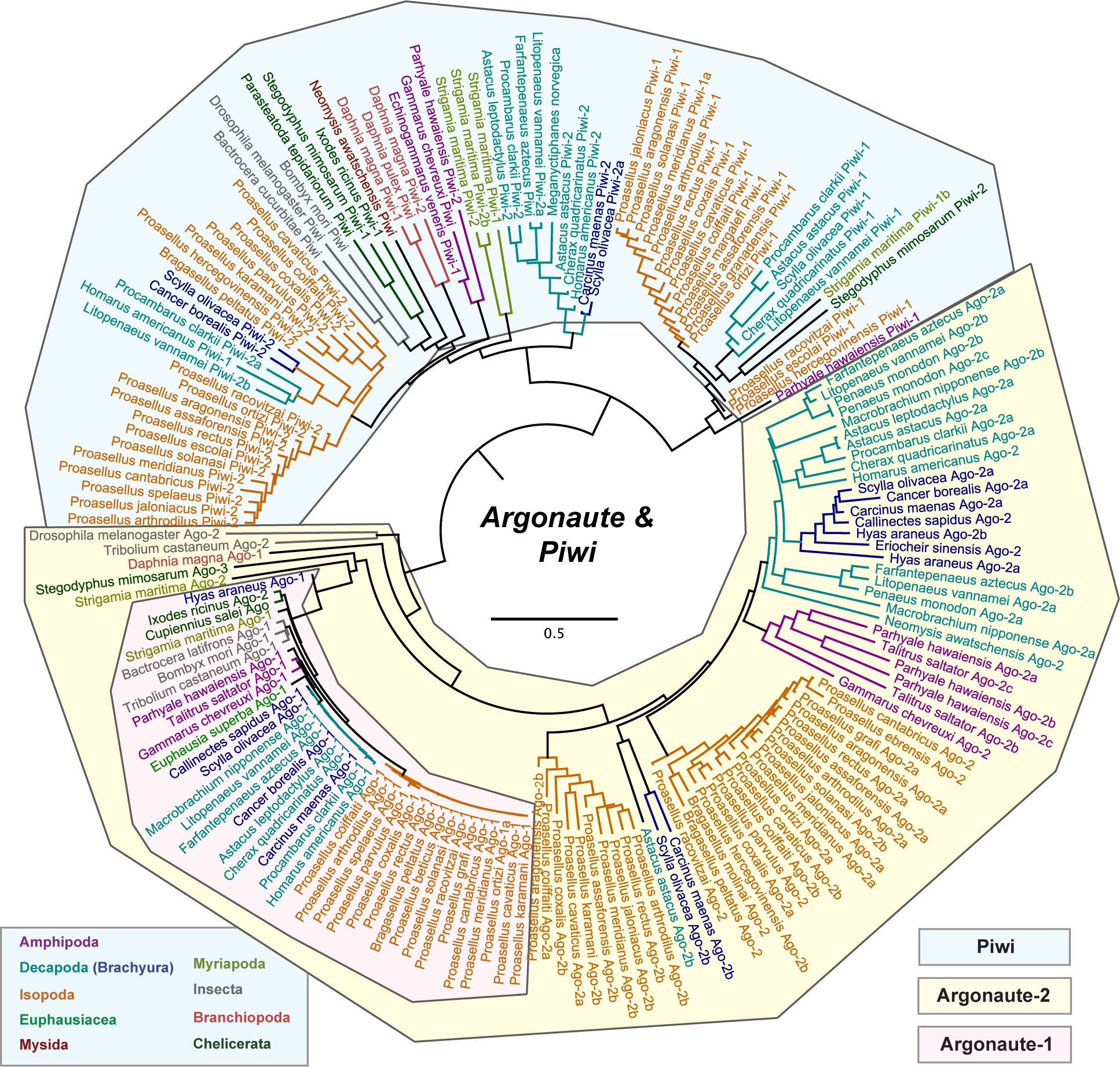
*Argonaute* and *Piwi* gene families in malacostracans. Phylogenetic tree of the *Argonaute* and *Piwi* family is constructed using the maximum-likelihood method from an amino acid multiple sequence alignment. Seven main groups of SOCS proteins are identified in malacostracans. Taxa labels are depicted as their respective colour codes. Bootstrap support values (n=1000) for all trees can be found in supplementary figure 8. Scale bar represents substitution per site.

